# Experimental evidence pointing to rain as a reservoir of tomato phyllosphere microbiota

**DOI:** 10.1101/2021.04.08.438997

**Authors:** Marco E. Mechan-Llontop, Long Tian, Parul Sharma, Logan Heflin, Vivian Bernal-Galeano, David C. Haak, Christopher R. Clarke, Boris A. Vinatzer

## Abstract

Plant microbiota play essential roles in plant health and crop productivity. Comparisons of community composition have suggested seeds, soil, and the atmosphere as reservoirs of phyllosphere microbiota. After finding that leaves of tomato (*Solanum lycopersicum*) plants exposed to rain carried a higher microbial population size than leaves of tomato plants not exposed to rain, we experimentally tested the hypothesis that rain is a so far neglected reservoir of phyllosphere microbiota. Rain microbiota were thus compared with phyllosphere microbiota of tomato plants either treated with concentrated rain microbiota, filter-sterilized rain, or sterile water. Based on 16S rRNA amplicon sequencing, one-hundred and four operational taxonomic units (OTUs) significantly increased in relative abundance after inoculation with concentrated rain microbiota but no OTU significantly increased after treatment with either sterile water or filter-sterilized rain. Some of the genera to which these 104 OTUs belonged were also found at higher relative abundance on tomatoes exposed to rain outdoors than on tomatoes grown protected from rain in a commercial greenhouse. Taken together, these results point to precipitation as a reservoir of phyllosphere microbiota and show the potential of controlled experiments to investigate the role of different reservoirs in the assembly of phyllosphere microbiota.

## INTRODUCTION

Microbial communities associated with plants, often referred to as plant-associated microbiota and as constituents of the plant microbiome, influence a remarkable number of processes of plant biology and affect plant health and crop yield (Goh et al., 2013, Badri et al., 2013, Hacquard et al., 2015, Lu et al., 2018, Torres-Cortés et al., 2018, Durán et al., 2018, Ritpitakphong et al., 2016, Berg & Koskella, 2018). The phyllosphere, considered here as the plant compartment that extends from the outside to the inside of the leaf (Morris, 2002, Vacher et al., 2016), harbors a high diversity of microorganisms, with bacteria being the most abundant domain (Lindow & Brandl, 2003, Vorholt, 2012).

Phyllosphere microbiota are exposed to fluctuating environmental stresses, including changes in UV exposure, temperature, water availability, osmotic stress, and humidity (Hirano & Upper, 2000, Jacobs & Sundin, 2001, Lindow & Brandl, 2003, Vacher et al., 2016). The core bacterial phyla that can withstand these environmental stressors include Proteobacteria, Actinobacteria, Bacteroidetes, and Firmicutes (Vorholt, 2012, Bulgarelli et al., 2013). However, at lower taxonomic ranks, the phyllosphere microbiome greatly varies with changing biotic and abiotic factors (Lindemann & Upper, 1985, Rastogi et al., 2012, Copeland et al., 2015, Wagner et al., 2016).

Culture-independent 16S rRNA amplicon analyses have expanded our knowledge of the composition of the phyllosphere microbiome of several plant species (Knief et al., 2012, Williams et al., 2013, Kembel et al., 2014, Copeland et al., 2015, Grady et al., 2019), including tomato (*Solanum lycopersicum*) (Ottesen et al., 2013, Ottesen et al., 2016). The bacterial genera *Pseudomonas*, *Erwinia*, *Sphingomonas, Janthinobacterium, Curtobacterium, Agrobacterium, Stenotrophomonas, Aurantimonas, Thermomonas, Buchnera, Enterococcus*, *Rubrobacter*, *Methylobacterium*, *Deinococcus*, and *Acidovorax* have all been observed to be associated with the phyllosphere of tomatoes grown in the field (Ottesen et al., 2013, Ottesen et al., 2016, Toju et al., 2019).

Microbes constantly cycle across interconnected habitats maintaining healthy ecosystems (van Bruggen et al., 2019). Importantly for this study, it has been shown that plants represent an important source of microbes that are released as aerosols into the atmosphere (Lindemann et al., 1982, Constantinidou, 1990, Lighthart & Shaffer, 1995, Šantl-Temkiv et al., 2013, Bowers et al., 2011, Vaïtilingom et al., 2012). The atmosphere then serves as a vehicle for microbial dispersal not only locally, but also globally (Bovallius et al., 1978, Brown & Hovmøller, 2002, Schmale & Ross, 2015) and air-borne microbial communities are deposited back to earth surfaces as precipitation. The atmosphere thus represents a crucial route for the dissemination of beneficial and pathogenic species (Polymenakou, 2012, Monteil et al., 2014, Monteil et al., 2016). There is even some evidence that air-borne microbes contribute to the formation of precipitation itself (Christner et al., 2008, Morris et al., 2014, Amato et al., 2015, Amato et al., 2017, Failor et al., 2017). Proteobacteria followed by Bacteroidetes, Actinobacteria, and Firmicutes have been identified as the most common phyla both in the atmosphere (Peter et al., 2014, Hiraoka et al., 2017, Cáliz et al., 2018) as in precipitation (Cáliz et al., 2018, Aho et al., 2020), notable largely the same as the core phyla of the phyllosphere microbiome mentioned earlier.

Microbial community assembly is largely influenced by deterministic (selection) and stochastic (dispersal) processes that determine the complex structure and function of microbiomes (Powell et al., 2015, Zhou & Ning, 2017, Graham & Stegen, 2017). This assembly process has been extensively studied in roots, which recruit microbial communities from the surrounding soil (Fitzpatrick et al., 2018, Pérez-Jaramillo et al., 2019). However, sources that drive the phyllosphere microbiome assembly are still under debate. It has been suggested that soil is the major reservoir of the phyllosphere microbiome (Zarraonaindia et al., 2015, Wagner et al., 2016, Grady et al., 2019). In fact, it has been observed that the leaf microbiome reflects soil bacterial diversity at an early stage of growth but significantly differs as plants grow and mature (Copeland et al., 2015). On the contrary, (Maignien et al., 2014, Ottesen et al., 2016) found evidence that dry deposition of air-borne microbes constitutes an important source of phyllosphere microbiota.

A few studies have also explored wet deposition of air-borne microbes in precipitation as a potential reservoir of phyllosphere microbiota. For example, (Morris et al., 2008) concluded that the plant pathogen *Pseudomonas syringae* is disseminated through the water cycle because of its ubiquitous presence in compartments of the water cycle and plants. Using whole genome sequencing, (Monteil et al., 2016) confirmed these conclusions by finding that *P. syringae* bacteria isolated from diseased cantaloupe plants and *P. syringae* isolates from rain, snow, and irrigation water were members of the same population. Recently, rain was also found to affect the overall composition of plant phyllosphere microbiota but it was not determined if rain-borne bacteria were the source of the observed shifts (Allard et al., 2020). Moreover, it is well known that fungal spores are released from plants, travel long distances through the atmosphere, and can be deposited back on plants with rain (Woo et al., 2018).

After finding that the phyllosphere of tomato plants exposed to rain contained a higher bacterial population size than the phyllosphere of tomato plants not exposed to rain, we hypothesized that rain-borne bacteria might contribute to the assembly of phyllosphere microbiota beyond pathogenic bacteria and fungi. Putative rain-borne tomato phyllosphere colonizers were identified in a series of controlled laboratory experiments and by comparing the composition of tomato phyllosphere microbiota of plants naturally exposed to rain with those not exposed to rain.

## MATERIALS AND METHODS

### Determination of the population size of tomato phyllosphere microbiota

Store-bought tomato seeds of the cultivar ‘Rio Grande’ were germinated in autoclaved soil (60 min/ fast cycle). For growth under laboratory conditions, plants were kept for 4 weeks on shelves under 14 h of light and 10 h of dark. For exposure to outdoor conditions, 4-week-old tomato plants were transplanted into large plastic pots. Pots were then either transported to a research farm and placed on gravel near a maintained lawn (Kentland Farm, Blacksburg, VA, USA) or put on the flat roof of the 3-story Latham Hall research building at Virginia Tech (Blacksburg, VA, USA) several meters above and away from soil and vegetation.

### Culture-dependent analysis of phyllosphere microbiota

Leaf disks were aseptically collected with a #1 cork borer and placed in a tube containing 200 µL of sterile 10 mM MgSO_4_ solution and 3 glass beads. Tubes were placed in a mini bead beater (Biospec Products, Inc., Bartlesville, OK, USA) and shaken for 2 min to release bacterial cells. Serial dilutions were plated on R2A plates supplemented with cycloheximide to inhibit fungal growth. Plates were incubated at room temperature and colony-forming units were counted 4 days later.

### Rain collection

Rain was collected as described in (Failor et al., 2017). In short, autoclavable bags were wrapped in aluminum foil and autoclaved for 40 min/fast cycle. Trash cans were arranged away from structures on the roof of the Latham Hall research building. Surfaces of containers were sprayed with 75% ethanol to prevent contamination. Sterile bags were placed in the cans and the lid placed back on top until the beginning of a rainfall event, at which point they were removed. The lids of three cans were then removed but one can was kept closed during the precipitation event as a negative control. After the end of precipitation events, 1 liter (L) of sterile water was poured into the negative control can, simulating the precipitation event. After rainfall events ended, bags containing rain water were removed and placed at 4^°^C until processing.

For DNA extraction, three L of rainwater were vacuum filtrated (reusable filter holders from Thermo Scientific Nalgene, USA) through a 0.2 µm pore filter membrane (Supor^®^ 200 PES membrane Disc Filter, PALL, USA). Filters were removed using sterile tweezers, placed into a 15 mL Eppendorf tube and stored at −80°C until processing.

### Rain as bacterial inoculum and plant treatments

Tomato plants were grown in the laboratory for four weeks as described above for the culture-dependent analysis of phyllosphere microbiota. Two L of rainwater were vacuum-filtered as described under rain collection. To concentrate the bacterial microbiota present in rain hundred fold, the filter membranes were incubated for 10 min with shaking in 20 mL of sterile water, which was then used as inoculum (referred to as concentrated rain microbiota or 100X-rain from now on). The rain that passed through the filter was used as bacterial-free inoculum (referred to as filtered rain or filter-sterilized rain). Autoclaved double-distilled water was used as sterile water treatment. Groups of four plants placed together into 13 inch x 16 inch plastic bags were sprayed until run off. The bags were left open for two hours to let the plants dry before the day 0 time point DNA extraction from two of the four tomato plants. Bags were then closed for 2 days to create a high humidity environment favorable to plant colonization after which they were kept open for 5 days until the day 7 time point DNA extraction from the remaining two tomato plants.

### Growth of outdoor and greenhouse tomatoes

Tomato plants were purchased at Home Depot at the beginning of June 2019 and transplanted into pots and placed on the flat roof of the Latham Hall building on the Virginia tech campus by setting them on a large, clean sheet of plastic to reduce contamination from the roof floor. Plants were regularly waters without touching leaves. One set of plants were brought indoors before each rainfall event while the other set of plants were kept outside to be exposed to rainfall. Each rainfall event was collected and DNA extracted as described above and leaves from both sets of plants were harvested and DNA extracted as described below.

### Harvesting plants leaves

All plant leaves were removed and collected in a ziplock plastic bag. Sterile distilled water was added (300 mL) and samples were sonicated for 10 minutes using a 1510 BRANSON sonicator (Brandsonic, Mexico) (Ottesen et al., 2013). The leaf wash was vacuum-filtered on to the same kind of 0.22 µm pore filter described above to collect microbial cells dislodged from leaf tissue during sonication, as described above. Filter membranes were placed into a 15 mL Eppendorf tube and stored at −80^°^C until processing.

Plant leaves were also collected using the same procedure inside a commercial greenhouse where tomatoes were grown either hydroponically or in non-sterilized soil under an organic production regiment.

### DNA extraction

DNA extraction from all the 0.22 µm filter membranes was performed using the Power Water DNA isolation kit (Qiagen, USA) according to the manufacturer’s protocol with minor modifications. DNA concentration and quality were assessed by UV spectrophotometry (NanoDrop 1000, Thermo, USA) and visualized on a 1 % agarose gel.

### Library preparation and sequencing

For the 16S rRNA amplicon sequencing we used the barcoded primers a799wF (5’AMCVGGATTAGATACCCBG3’) and new1193R (5’ACGTCATCCCCACCTTCC3’). A 28 cycle PCR was performed using the HotStarTaq Plus Master Mix Kit (Qiagen, USA) under the following conditions: 94°C for 3 minutes, followed by 28 cycles of 94°C for 30 seconds, 53°C for 40 seconds and 72°C for 1 minute, after which a final elongation step at 72°C for 5 minutes was performed. After amplification, PCR products obtained from the various samples were mixed in equal concentrations and purified using Agencourt Ampure beads (Agencourt Bioscience Corporation, Beverly, MA, USA). All steps from PCR to paired-end (2 × 300 bp) amplicon sequencing on the Illumina MiSeq platform were performed at Molecular Research LP (MR DNA™, Shallowater, TX, USA).

For metagenomic sequencing using the Illumina platform, total DNA was sequenced using 150 bp paired-end reads on the HiSeq 4000 Illumina platform, Duke University Sequencing and Genomic Technologies Shared Resource, Durham, NC, USA. For metagenomic sequencing using the Nanopore platform, DNA libraries were prepared following the ‘1D Native barcoding genomic DNA protocols (SQK-LSK109 and EXP-NBD104) provided by Oxford Nanopore Technologies (ONT).

### Bioinformatic analysis

Raw 16S rRNA paired-end sequences were processed by Molecular Research LP (MR DNA™, Shallowater, TX, USA) as follows: 1) reads were joined together after q25 trimming of the ends and reoriented in the 5’-3’ direction, 2) barcodes and primer sequences were removed, and 3) sequences shorter than 200bp, sequences with ambiguous base calls, and sequences with homopolymer runs exceeding 6 bp were removed. Operational taxonomic units (OTU) were assigned using the open source Quantitative Insights into Microbial Ecology (QIIME) version 1.9.1bioinformatic pipeline (Caporaso et al., 2010), using the open-reference protocol at 97% sequence identity with UCLUST as the clustering tool and SILVA release 128 (Quast et al., 2013) as the database. All OTUs annotated as mitochondria, chloroplasts, cyanobacteria, unassigned, and OTUs with fewer than five reads were removed from the dataset.

The QIIME-generated output file in the Biological Observation Matrix (BIOM) format was used for downstream data analysis and visualization in R version 3.3 using the Vegan (Oksanen J., 2020), Phyloseq 1.26.1 and ggplot2 3.3.2 packages (McMurdie & Holmes, 2013, Hadley, 2009).

Samples were rarefied to the lowest sample depth to compute diversity analysis. The rain core microbiome analysis was performed with the microbiome R package (Lahti L., 2020) using a detection threshold of 0.1% and 100% as a prevalence threshold. Alpha diversity was assessed using observed OTUs, Shannon, and Simpson indices. Differences in alpha diversity were determined by pairwise Wilcoxon rank sum test with the Holm correction method. Beta diversity was analyzed based on unweighted UniFrac distance, weighted UniFrac distance, and Bray-Curtis dissimilarity. Differences in beta diversity were determined using PERMANOVA as implemented in adonis2 (from vegan 2.5-6 using models with 999 permutations, adonis2(dist.matrix ∼Treatment*TimePoint + DateExperiment, permutations = 999). Dissimilarity matrices were visualized using the Principal Coordinates Analysis (PCoA) ordination method as implemented in the Phyloseq package. DESeq2 (Love et al., 2014) was used to identify OTUs that were differentially abundant across treatment groups and time points. OTUs were filtered using a False Discovery Rate (FDR) cutoff of 0.01.

For metagenomic data analysis, raw 150 bp paired-end reads from Illumina were processed to remove short and low quality reads using Trimmomatic (version 0.38) (Bolger et al., 2014). Reads with an average per base quality below 30 and read length below 150bp were filtered out. Filtered reads were then classified taxonomically using Centrifuge (version 1.0.4) (Kim et al., 2016) and Sourmash version 2.0.0 (Brown, 2016) only retaining species that were identified by both classifiers.

For Oxford Nanopore Technologies sequencing, the fast5 files containing the raw reads obtained from the MinION sequencer were base-called using Guppy (v3.3.2). The ONT workflow What is in my pot (WIMP v2019.7.9) was used for bacterial identification and a classification (Juul et al., 2015).

## RESULTS

### The bacterial population size on tomato plants exposed to rain is larger than that of tomato plants not exposed to rain

Our investigation into the role of rain in shaping the phyllosphere microbiome started by comparing the bacterial population size on tomato plants grown indoors under controlled conditions with that of plants grown outside exposed to environmental disturbances including rainfall. We observed that tomato plants grown over four weeks outdoors in plastic pots at the Virginia Tech Kentland research farm harbored bacterial populations of significantly larger size compared to plants grown under laboratory conditions for the same period (Figure 1A). Interestingly, even plants grown on the roof of a campus research building, which minimized microbial dispersal from soil and plants compared to the farm environment, had bacterial populations that were significantly larger compared to those of plants grown indoors (Figure 1B). These results suggest that air-borne and rain-borne bacteria through dry and/or wet deposition had colonized the tomato plants grown outside.

**Figure 1.**
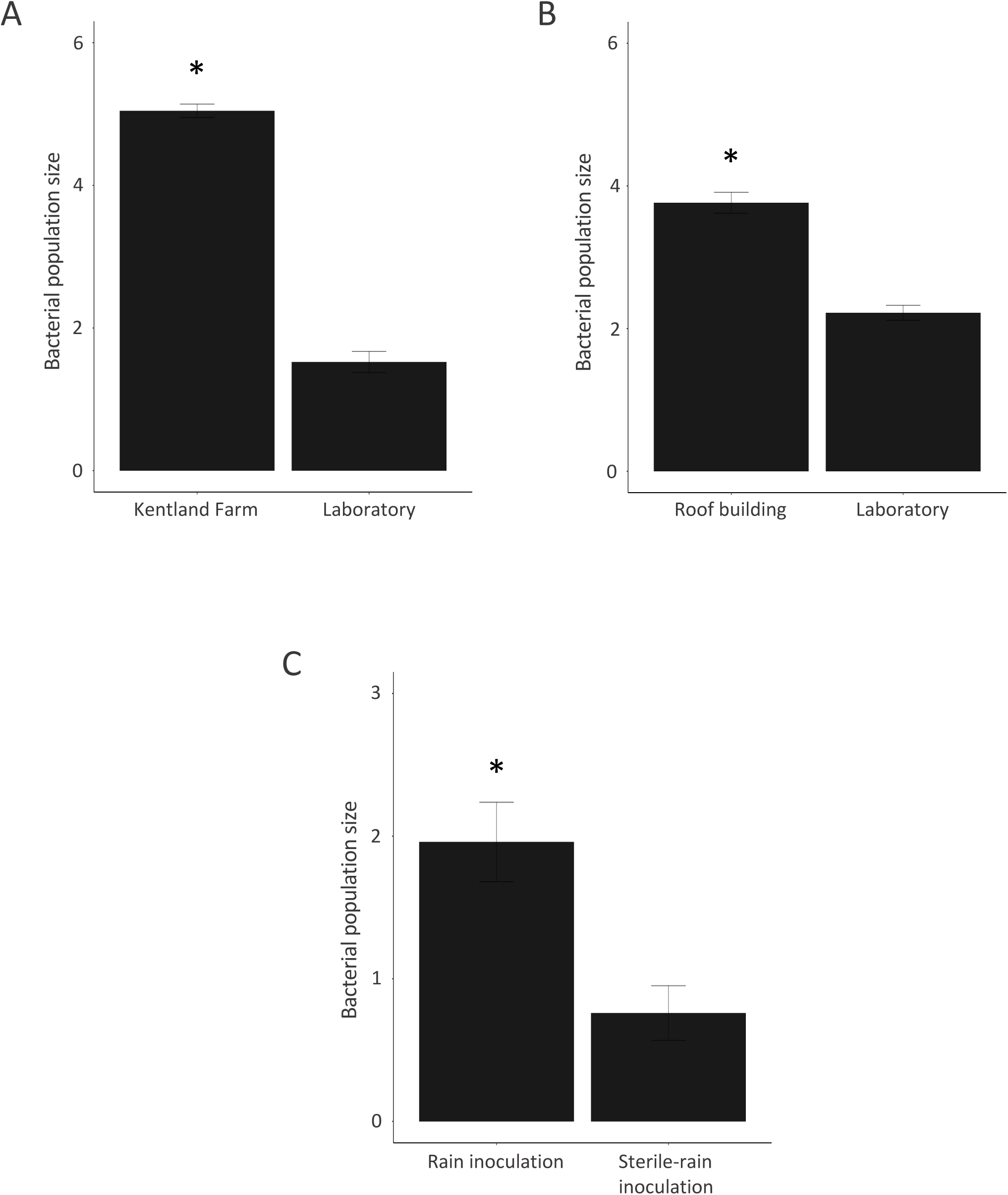
Bacterial population size in the phyllosphere of tomato plants A) grown exposed to rain in plastic pots at the Virginia Tech Kentland Farm compared to plants grown inside the laboratory B) grown on the roof of the Latham Hall research building exposed to rain compared to plants grown under laboratory conditions, and C) grown inside the laboratory seven days after being either treated with rain or autoclaved rain. T-test, P < 0.001.

To test the effect of rain on the bacterial population size in the tomato phyllosphere under controlled conditions, we collected rain and used it as an inoculum to treat 4-week old tomato plants that had been grown under laboratory conditions. Seven days after plants were treated with rain, they carried a significantly larger bacterial population count compared to plants that had been treated with autoclaved rain (Figure 1C). This result suggested that at least some rain-borne bacteria are able to colonize plant leaves efficiently and may thus impact bacterial population composition in the phyllosphere.

### Rain-borne microbiota in Blacksburg, Virginia, are highly variable

As a first step towards identifying which bacterial taxa present in rainfall may efficiently colonize the tomato phyllosphere, we characterized the bacterial diversity associated with nine rainfall events during 2015 and 2016. Rainfall was collected in sterile plastic bags on the same roof of the research building previously used to grow tomatoes, DNA was extracted, and the 16S rRNA gene was amplified and sequenced (Table 1). In total, 1,186,365 short reads were obtained. After 97% OTU clustering and removal of all non-bacterial and unassigned reads, a total of 892,142 reads remained. All samples were rarefied to 6419 reads per sample and 5958 OTUs overall were identified. Rarefaction curves (Supplementary Figure 1) show that this is an underestimate of the total number of OTUs since not all samples were sequenced to saturation. The number of OTUs per rarefied sample ranged from 541 (June 2019) to 1782 (April 2016).

**Table 1.**
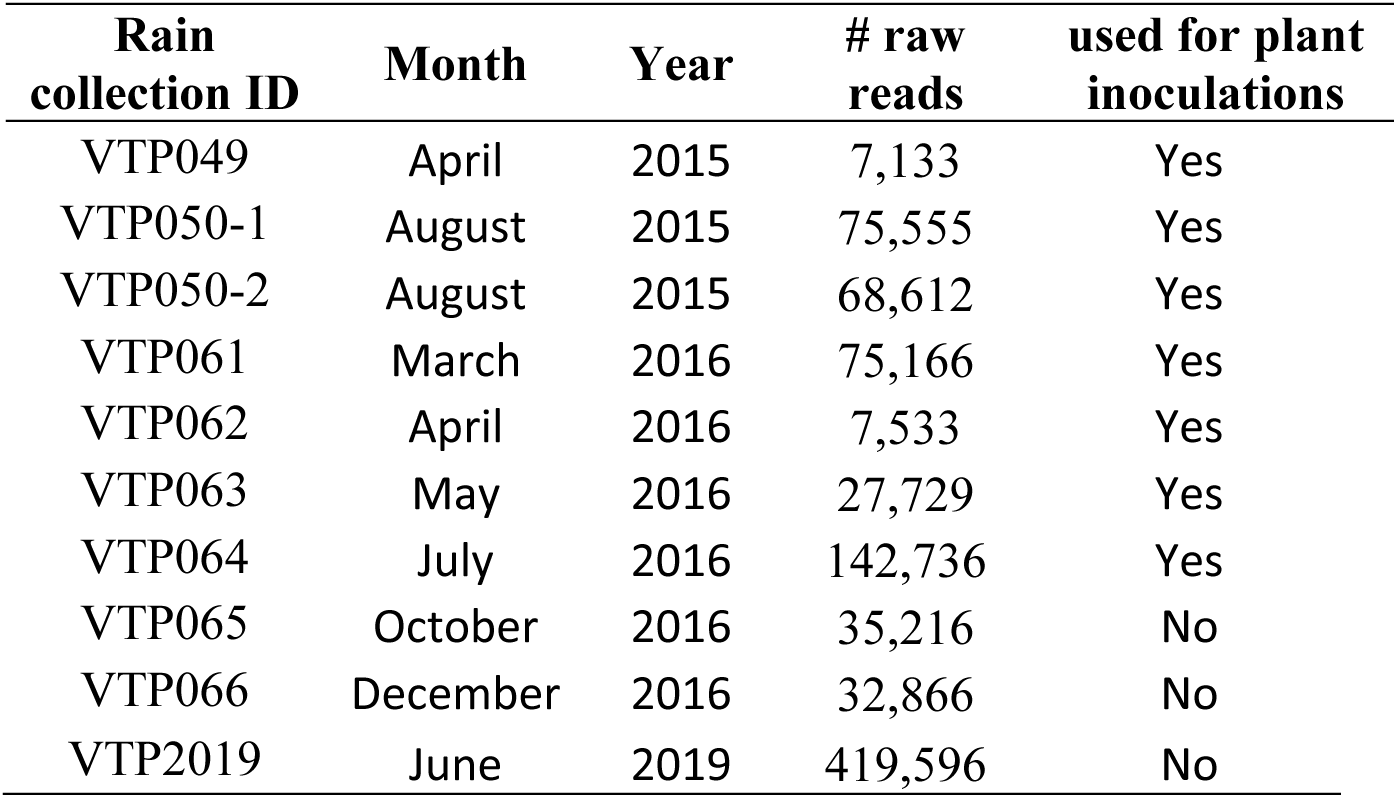
Metadata for analyzed rain samples.

| **Rain**<br>**collection ID** | **Month** | **Year** | **# raw**<br>**reads** | **used for plant**<br>**inoculations** |
| --- | --- | --- | --- | --- |
| VTP049 | April | 2015 | 7,133 | Yes |
| VTP050-1 | August | 2015 | 75,555 | Yes |
| VTP050-2 | August | 2015 | 68,612 | Yes |
| VTP061 | March | 2016 | 75,166 | Yes |
| VTP062 | April | 2016 | 7,533 | Yes |
| VTP063 | May | 2016 | 27,729 | Yes |
| VTP064 | July | 2016 | 142,736 | Yes |
| VTP065 | October | 2016 | 35,216 | No |
| VTP066 | December | 2016 | 32,866 | No |
| VTP2019 | June | 2019 | 419,596 | No |

Taxonomic diversity analysis revealed Proteobacteria, Actinobacteria, Bacteroidetes, Acidobacteria, and Firmicutes to be the dominant taxa at the phylum level. At the class level, Alphaproteobacteria followed by Gammaproteobacteria, Actinobacteria, Acidobacteria, and Bacilli were most abundant. However, there were considerable differences among samples. For example, Chlamydiae represented the most abundant taxon in the rain sample collected in August 2015 at 36% relative abundance, represented only 0.4% in the sample collected in July 2016, and were less than 0.1% in all other samples. Deltaproteobacteria were only present in August 2015, May 2016 and July 2016 at 12.79%, 1.42%, and 17.03% relative abundance, respectively. Actinobacteria were most abundant in rain samples collected in April 2016 (25.56%) and December 2016 (16.29%) while they only represented 2.18% of the October 2016 sample. Gammaproteobacteria represented 80% of relative abundance in June 2019 but only averaged 28.41% in the other samples (Figure 2A).

**Figure 2.**
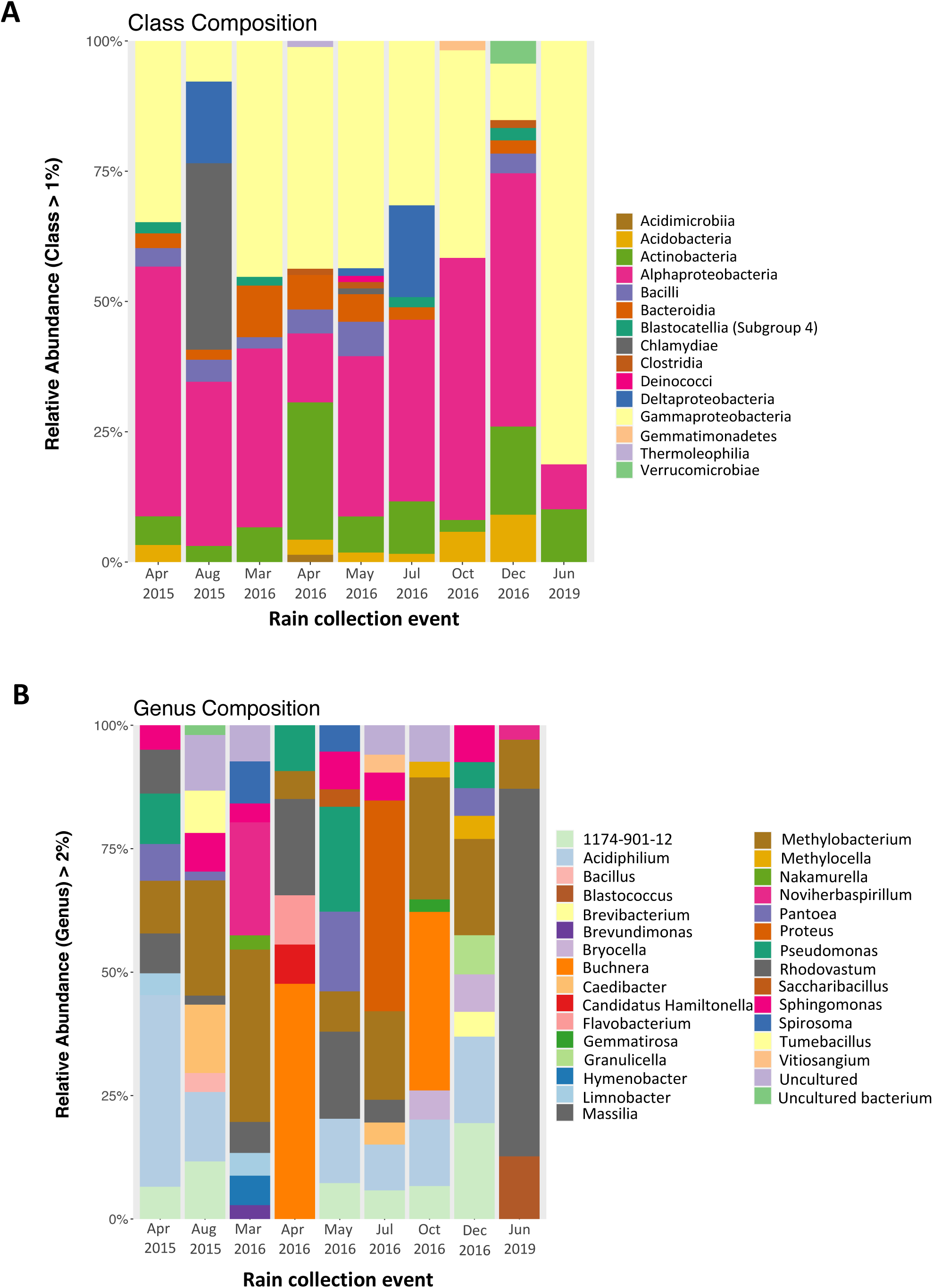
Relative abundance (RA) of bacterial taxa in rainfall collected in Blacksburg, VA, on 9 different days in 2015, 2016, and 2019. Samples were rarefied to 6,419 sequences. A) RA at the class level (only classes with a RA above 1% are shown), B) RA at the genus level (only generated with RA above 2% are shown). For August 15, results are based on two technical replicates (see also Table 1).

Even more dramatic differences in relative abundance between samples were observed at the genus level (Figure 2B). However, seventeen OTUs were identified across all nine samples when setting a detection threshold of 0.1%. These OTUs belong to the following genera (listed in order of decreasing relative abundance): *Acidiphilium*, *Bryocella*, Beijerinckiaceae 1174-901-12, *Methylobacterium, Massilia*, Burkholderiaceae DQ787673.1.1527, *Pantoea*, *Pseudomonas* and *Sphingomonas* (Supplementary Figure 2).

### Concentrated rain microbiota, filter-sterilized rain, and sterile water all affect the phyllosphere of lab-grown tomato plants

Next, we decided to determine if inoculation with rain microbiota would not only increase the size of the tomato phyllosphere microbiota as seen in Figure 1 but also change its composition because of colonization of tomato leaves by rain-borne bacteria. In six independent experiments, 100-fold concentrated rain microbiota (100X-rain) derived from six out of the nine collected rainfall events described above (April 2015, August 2015, March 2016, April 2016, May 2016, July 2016) were used to inoculate tomato plants. In parallel, a separate set of tomato plants were treated each time with filter-sterilized rain obtained from the concentration step above. For each of the 2016 experiments, additional tomato plants were treated with sterile water. 16S rRNA amplicons were prepared and sequenced from DNA extracted from leaf washes of tomato plants two hours after treatments (day-0) and seven days later (day-7). For the March 2016 and May 2016 experiments enough rain and enough plants were available so that each treatment was done in duplicate. For the April 2016 experiment, two day-7 100X-rain samples were taken. For the other three experiments, only one sample per treatment and time point was processed.

In total, 3,291,016 reads were obtained from 45 phyllosphere samples (Supplementary Table 1). After 97% OTU clustering and removing all non-bacterial and unassigned reads, a total of 3,118,320 reads remained in the data set. Rarefaction curves show that most of the samples were deeply sequenced (Supplementary Figure 1).

After subsampling to 6,670 reads per sample, we identified a total of 9,923 OTUs and measured the alpha diversity based on the total number of observed species and by Shannon and Simpson diversity indices. Figure 3 shows the alpha diversity values for rain compared to treated plants at day 0 and day 7. Although alpha diversity of rain microbiota was highly variable, the number of observed OTUs in rain was significantly higher than day-7 samples for all three treatments (p-values of 0.020, 0.050, and 0.028 respectively as determined by the pairwise Wilcoxon rank sum test with the Holm correction method) (Supplementary Table 2). Also, the number of observed OTUs significantly decreased in plants treated with 100X-rain from day 0 to day 7 (p-value 0.027). Comparing Shannon’s index, we observed a depletion in richness from day 0 to day 7 for plants treated with 100X-rain (p-value 0.049) as well as for plants treated with filtered-rain (p-value 0.023). No significant differences in alpha diversity were observed between day 0 and day 7 for plants treated with sterile-water.

**Figure 3.**
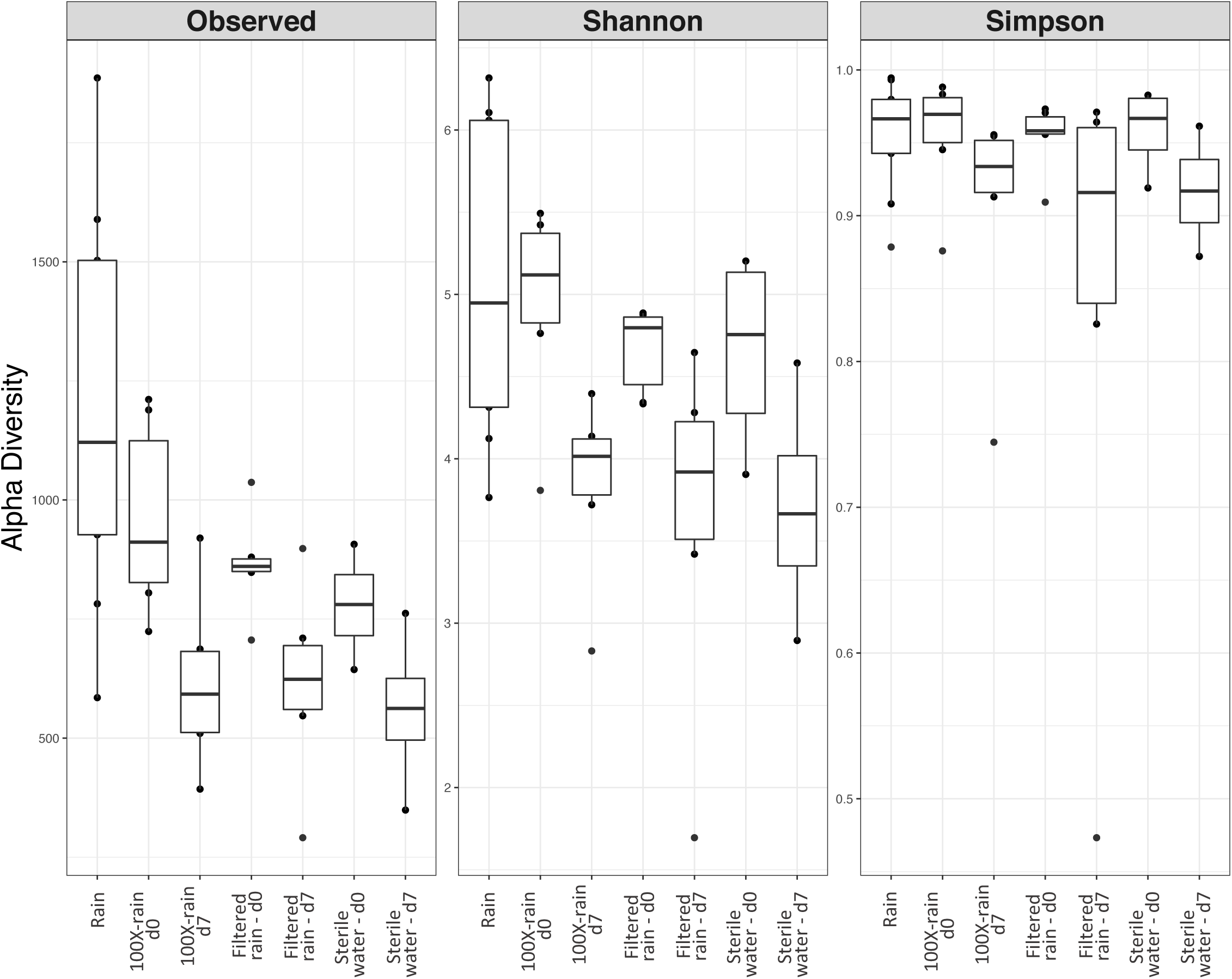
Alpha diversity measurements for rain compared to treated plants at day-0 vs day-7. Three measures of alpha diversity (observed OTUs, Shannon diversity index, and Simpson diversity index) were used. For rain, only samples used for plant inoculations were included.

Principal coordinate analysis (PCoA) derived from weighted and unweighted UniFrac distance metrics and from Bray-Curtis dissimilarity (Figure 4) revealed that most rain samples clustered together while phyllosphere samples did not. While PCoA derived from weighted Unifrac distances only separated phyllosphere samples along the second coordinate, PCoA derived from unweighted Unifrac distances and Bray-Curtis dissimilarity separated phyllosphere samples along both axes. Therefore, bacterial communities on lab-grown tomato plants were even more dissimilar from each other than the dissimilarity between rain communities. A permutational multivariate analysis of variance (PERMANOVA) for Bray-Curtis dissimilarity revealed that date of experiment, day of sampling (day 0 versus day 7), and treatment (100X-rain, filter-sterilized rain, and sterile water) were all significantly associated with bacterial community composition. (Supplementary Table 3). In regard to the date of experiment, it can be seen in panels 4B and 4C how samples clearly cluster by date of experiment along the X axis. Since this clustering is independent of treatment and day of sampling, the starting tomato phyllosphere population appears to have been different between experiment dates. Secondly, day 7 samples differed from day 0 samples on the Y axis independently of which treatment was applied revealing that inoculation and incubation under high humidity of plants by itself shifted the composition of the tomato phyllosphere population. Thirdly, although treatment was significant, there was interaction between treatment and date of experiment. Therefore, the effect of 100X-rain was not significantly different compared to the effect of filter-sterilized rain or sterile water. In other words, beta diversity analysis was unable to reveal if rain-borne bacteria present in the 100X-rain treatments affected the tomato phyllosphere community, possibly because of the differences in the composition of the microbial rain and phyllosphere communities between experiments.

**Figure 4.**
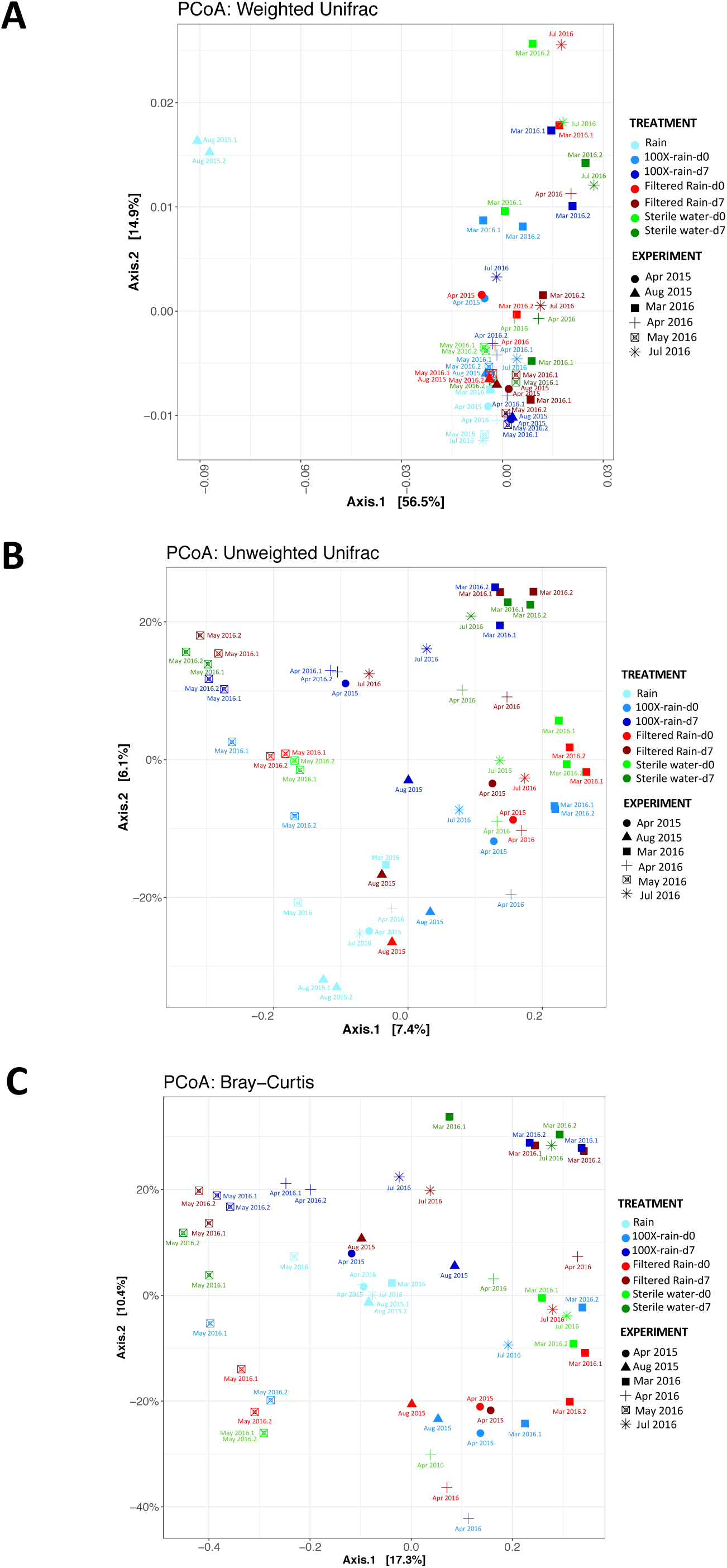
Principal coordinates analysis (PCoA) derived from weighted UniFrac distances (A), unweighted UniFrac distances (B), and the dissimilarity matrix of Bray-Curtis (C). Only rain samples used for plant inoculation were included. For experiments for which replicates were available, the replicate samples are labeled as “monthyear.1” and “monthyear.2”.

After finding that beta diversity analysis was inconclusive, we decided to compare the actual taxonomic diversity between samples. We observed an enrichment in Proteobacteria in the tomato phyllosphere on day 7 compared to the day 0 regardless of treatment. In contrast, relative abundance of Actinobacteria and Firmicutes was dramatically reduced 7 days post-treatment in all plants treated with 100X-rain but not after treatment with filter-sterilized rain or sterile water (Figure 5). At the class level, Gammaproteobacteria were significantly enriched in all tomato phyllosphere day 7 samples. Actinobacteria and Bacilli were reduced on day 7 in plants treated with 100X rain compared with filter-sterilized rain or sterile water. Unexpectedly, no consistent increase of any taxon at either phylum, class, or genus level was observed from day 0 to day 7 for plants treated with 100X-rain alone.

**Figure 5.**
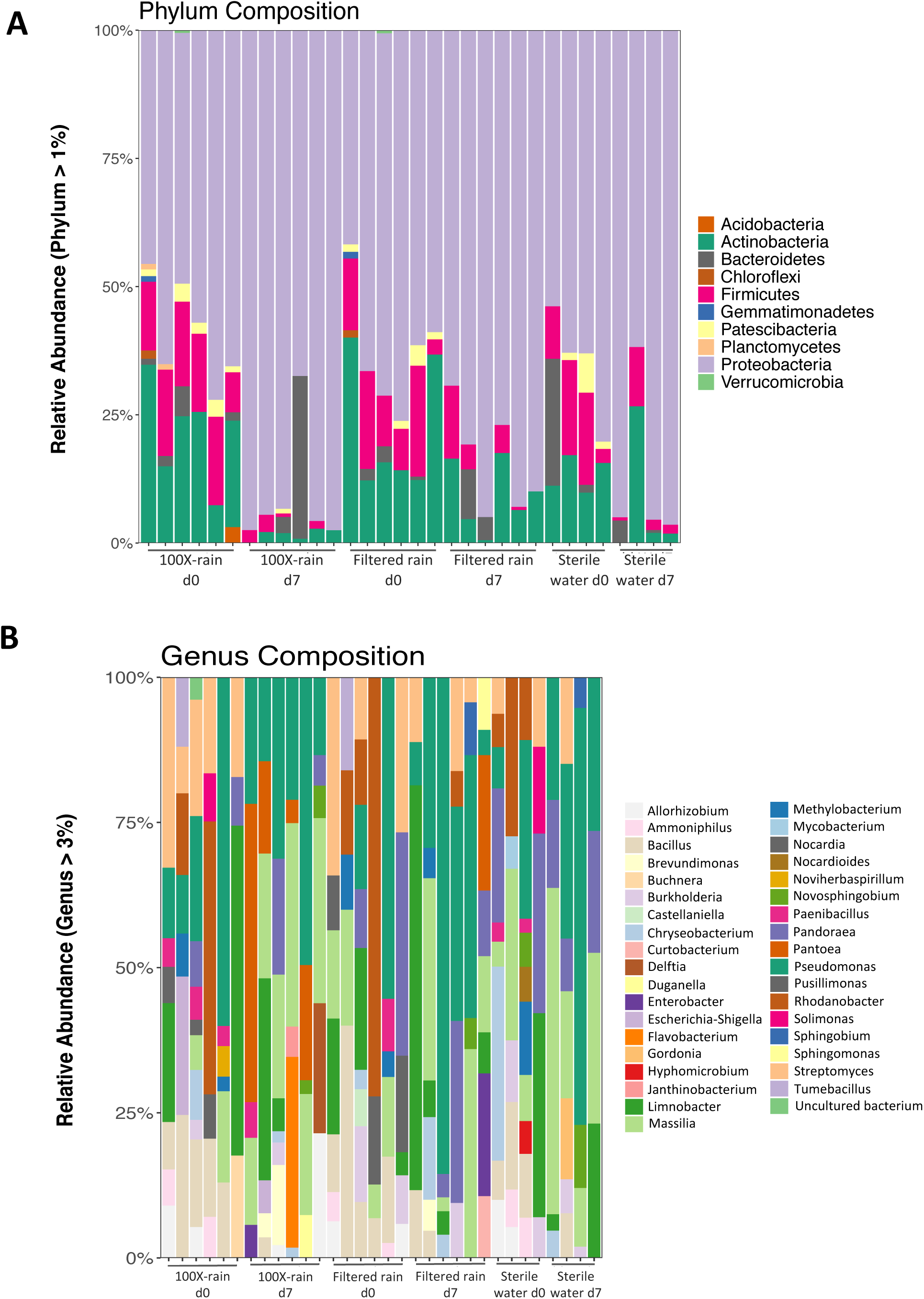
Relative abundance (RA) of bacterial taxa of plants treated with either 100X-rain, filtered rain, or sterile water, at day-0 and day-7. Experiments are listed by dates from left to right (April ‘15, August ‘15, March ‘16, April ‘16, May ‘16, and July ‘16). A) RA at the phylum level (abundance > 1%), B). RA at the genus level (abundance > 3%). For experiments for which replicates were available relative abundance is based on all replicates.

### One-hundred and four rain-borne OTUs increased significantly in relative abundance in the tomato phyllosphere exclusively after treatment with concentrated rain microbiota

Since we did not find any taxon between genus and phylum level that exclusively increased from day 0 to day 7 on tomato plants treated with 100X-rain, we wanted to determine if we could find any individual rain-borne OTUs that did so. To do this we used DESeq2 (Love et al., 2014) as it is relatively robust to small and unequal sample sizes as in the present study..

First, we directly compared day-0 and day-7 phyllosphere microbiota treated with 100X-rain and found that one-hundred and four OTUs (out of a total of 7,994 OTUs) significantly increased (Supplementary Table 4). These OTUs belonged to the genera *Massilia* (27 OTUs), *Pantoea* (18 OTUs), *Duganella* (13 OTUs)*, Pseudomonas* (11 OTUs), *Enterobacter* (5 OTUs), *Flavobacterium* (3 OTUs), *Janthinobacterium* (2 OTUs), and *Curtobacterium* (1 OTUs). 16 unknown OTUs from the *Burkholderiaceae* and 5 unknown OTUs from the *Enterobacteriaceae* families significantly increased in relative abundance as well (Figure 6A). Importantly, not a single OTU significantly differed in abundance between day 0 and day 7 on tomato plants treated with either filtered-rain or dd-water. This suggests that the OTUs which increased in relative abundance from day 0 to day 7 on tomatoes after treatment with 100X-rain originated from rain and were not members of the bacterial community present in the tomato phyllosphere prior to inoculation.

**Figure 6:**
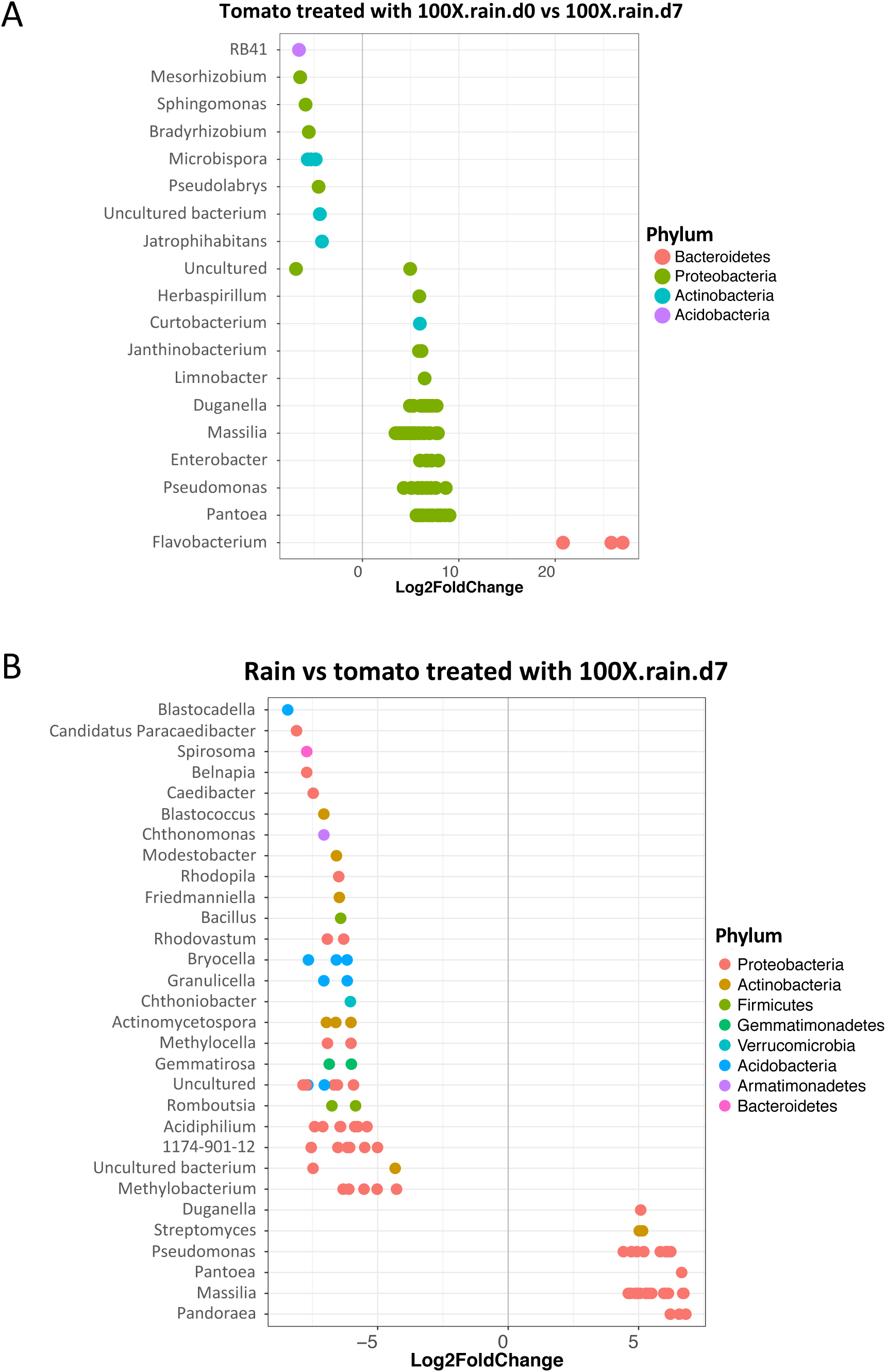
Differential abundance analysis at the level of Operational Taxonomic Unit (OTU) using DESeq2 (Love et al., 2014). The fold change is shown on the X axis and genera are listed on the Y axis. Each colored dot represents a separate OTU. A) Comparison of phyllosphere microbiota of plants treated with 100X-rain between day-0 and day-7, B) Comparison between rain microbiota and phyllosphere microbiota 7 days after treatment with the respective 100X-rain sample.

We then complemented the previous comparison with a slightly different analysis comparing the bacterial composition of rain microbiota with phyllosphere microbiota seven days after treatment with 100X-rain. We observed that thirty-five rain-borne OTUs out of a total of 5,958 OTUs had a significantly higher relative abundance on tomatoes at day-7 compared to their relative abundance in rain (Figure 6B). These OTUs (in order of decreasing differential abundance) mostly belonged to the same genera as the genera identified in the day-0 to day-7 comparison : *Massilia, Pseudomonas*, *Pandoraea*, *Streptomyces*, and *Pantoea*. The genus *Pantoea* was the genus that most consistently increased in relative abundance. In fact, *Pantoea* was detected in all rain collections (Figure 7A) and reached a high relative abundance (between 4% - 44%) in the tomato phyllosphere after 7 days each time its abundance in rain was above 1% (observed 4 out of 6 times). Genus-level abundance in rain and phyllosphere samples are also shown for *Flavobacterium, Janthinobacterium*, *Pseudomonas, Methylobacterium*, and *Massilia*, which all successfully colonized the tomato phyllosphere each time they were detected in rain (Figure 7B-E).

**Figure 7.**
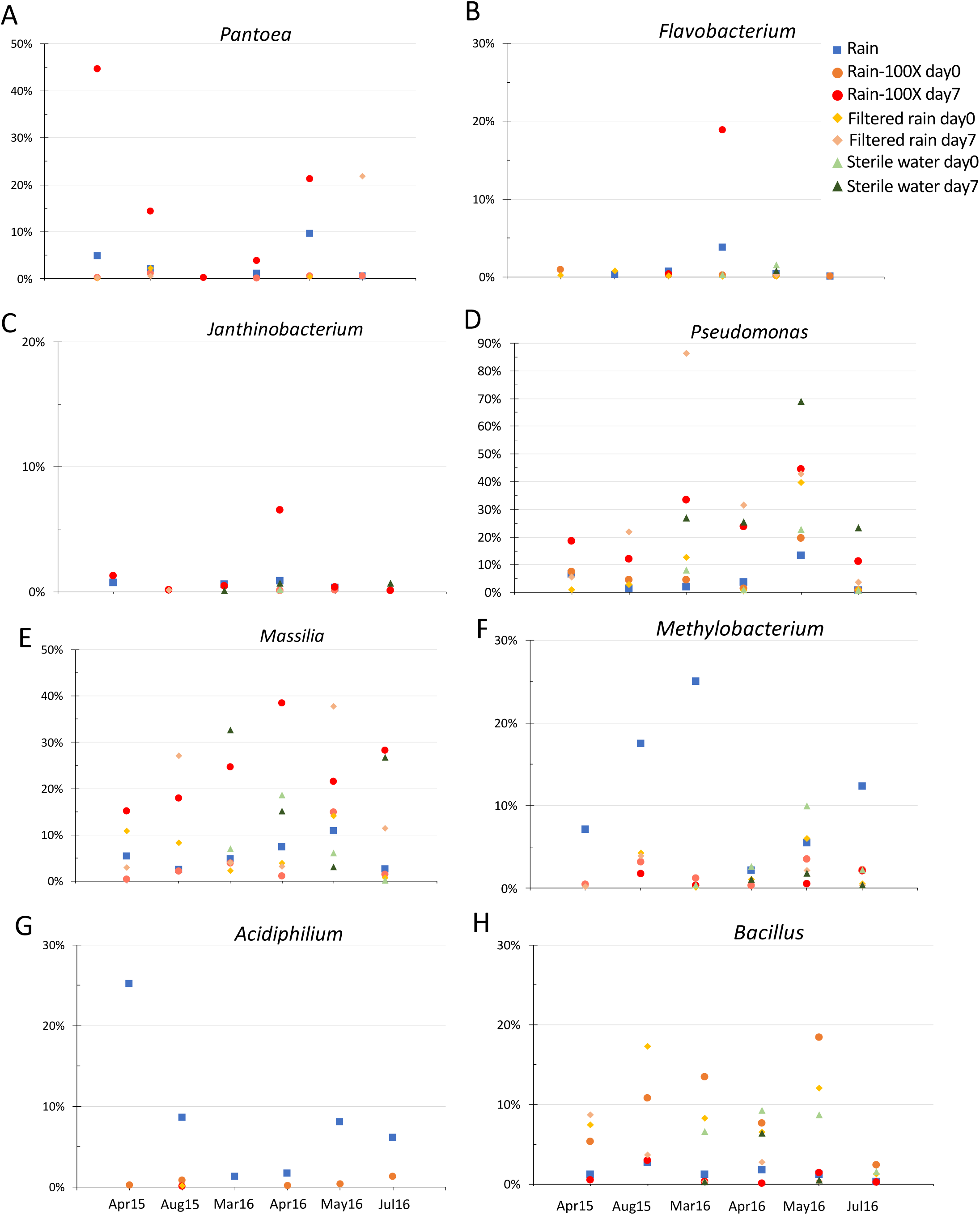
Relative abundance of a representative selection of rain-borne genera that either failed or succeeded in colonizing the tomato phyllosphere. Dates of experiments are listed on the X axis of panels G and H. Relative abundance is shown on the Y axis for rain, tomato plants on day-0 and day-7 after being treated with 100X-rain (Rain), filtered rain, or sterile water (see panel B for figure legend). A) *Pantoea*, B) *Flavobacterium*, C) *Janthinobacterium*, D) *Pseudomonas*, E) *Massilia*, F) *Methylobacterium*, G) *Acidiphilium*, and H) *Bacillus*.

In contrast, 61 OTUs found in rain samples significantly decreased in relative abundance by day 7 suggesting that these rain-borne taxa were definitely not able to colonize tomato leaves. These OTUs belonged to the following genera (list of the first 10 bacterial genera in order of decreasing differential abundance): *Acidiphilium*, Beijerinckiaceae-1174-901-12, *Bryocella, Actinomycetospora, Methylobacterium, Methylocella*, *Granulicella, Belnapia, Modestobacter*, and *Blastococcus.* The genus *Acidiphilium* best exemplifies this group of taxa. It was detected at high relative abundance in most rain samples, it was observed on plants treated with 100X-rain on day 0, but it was never found on plants treated with 100X-rain on day 7 (Figure 7G). This clearly shows that *Acidiphilium* is a common rain-borne bacterial genus that does not include any members able to colonize the tomato phyllosphere.

Several other pairwise comparisons were made to gain additional insights into differences in composition between microbiota at the OTU level. For example, we determined which OTUs were present in significantly higher abundance in rain compared to tomatoes treated with sterile water on day 0 to identify OTUs that are commonly present in rain but present in low abundance (or not at all) on laboratory-grown tomato plants (Supplementary Figure 3A). One hundred seventy-four such OTUs (out of a total of 5,958 OTUs present in rain) were identified. Most of these OTUs belonged to the following genera (list of the first 10 bacterial genera in order of decreasing differential abundance in rain): *Caedibacter*, *Tumebacillus*, *Acidiphilium*, *Methylobacterium, Rhodovastum, Belnapia, Sphingomonas, Methylocella*, *Bacillus and Pantoea*.

On the other hand, 109 OTUs had significantly higher relative abundance in laboratory-grown tomatoes (tomatoes at day 0 after being treated with sterile water) than in rain (Supplementary Figure 3A). These taxa are thus common inhabitants of tomatoes grown under our laboratory conditions in the absence of rain. They mostly belonged to the following genera (list of the first 10 bacterial genera in order of decreasing differential abundance): *Hyphomicrobium*, *Rhodanobacter*, *Chryseobacterium*, *Burkholderia*, *Paenibacillus*, *Nocardioides, Pandoraea*, *Bacillus*, *Novosphingobium*, and *Streptomyces*.

One genus that did not fall into any of the above categories was the genus *Bacillus*: *Bacillus* OTUs were observed at high relative abundance in rain samples as well as in the tomato phyllosphere at day 0 independent of treatment and on day 7 after sterile-rain and dd-water treatments but not after 100X-rain treatments (Figure 7H). This suggests that *Bacillus* OTUs were present in rain as well as on lab-grown tomatoes but were outcompeted by other rain-borne bacteria added with the 100X-rain treatments.

To more precisely identify the OTUs that represented the most efficient tomato colonizers, we used metagenome shotgun sequencing to re-sequence the microbiota associated with rain samples and with plants treated with 100X-rain on day 0 and day 7. The metagenome shotgun sequencing approach generated 260,035,170 short-reads. After quality control, 7,543,305 reads remained, of which 98.26% were identified as bacterial reads. Table 2 summarizes the results listing the bacterial species present in rain that numerically increased in abundance from day 0 to day 7 and ranked from high to low based on their relative abundance on day 7. Based on this analysis, rain-borne species *Pantoea vagans* and *Pantoea agglomerans* were the most effective tomato phyllosphere colonizers followed by *Pseudomonas citronellolis*, *Novosphingobium resinovorum*, an unnamed *Buttiauxella* species, *Erwinia gerundensis*, *Pseudomonas fluorescens*, *Cedecea neteri*, and an unnamed *Massilia* species. Additional *Pantoea*, *Massilia*, *Pseudomonas*, *Janthinobacterium*, and *Enterobacter* species ranked highly as well. The metagenomic analysis thus mostly confirmed and refined our 16S rRNA results of which rain-borne taxa are the most effective colonizers of the tomato phyllosphere.

**Table 2.**
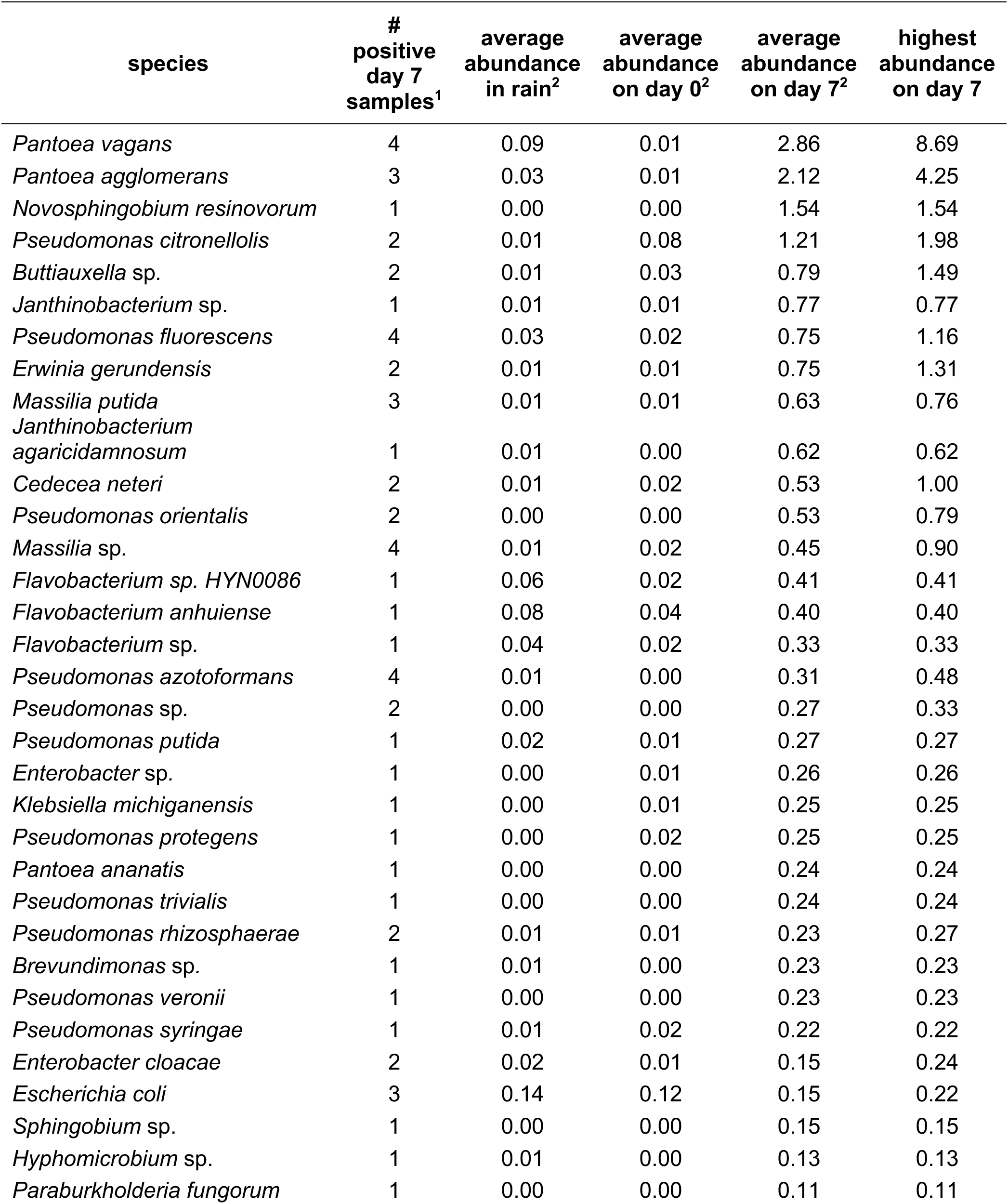
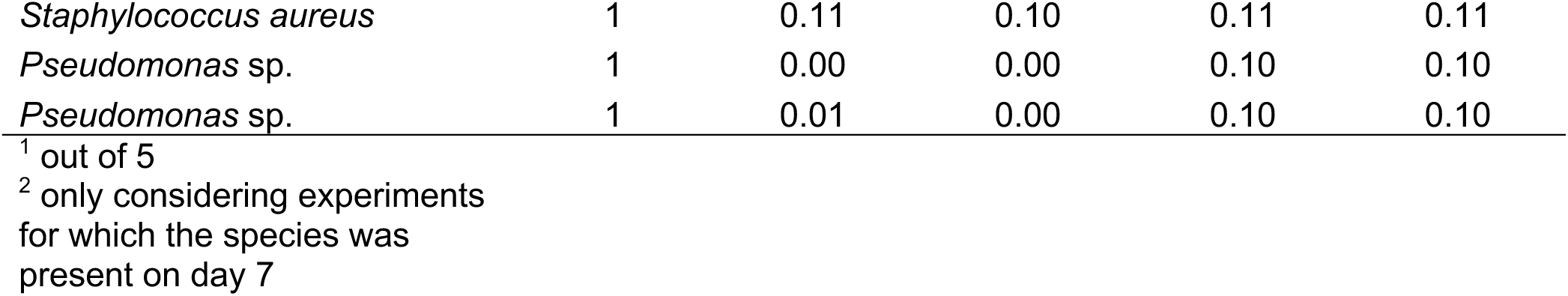
Species with highest relative abundance on tomato plants on day-7 after treatment with 100X-rain based on metagenomic sequencing.

### There are consistent differences between phyllosphere microbiota of tomato plants never exposed to rain and naturally exposed to rain outdoors

After identifying bacterial taxa present in rain that efficiently colonized tomato leaves under laboratory conditions, we tested the hypothesis that these taxa would be abundant in tomato plants grown outdoors in pots containing autoclaved soil on the roof of the same research building where we had collected rain samples earlier (7 samples) but be missing, or at least be underrepresented, in phyllosphere microbiota of greenhouse-grown tomatoes that had never been exposed to rain. This second set of plants included tomato plants grown in a hydroponic system (29 samples) and tomato plants grown in soil (18 samples), both in a commercial greenhouse (Supplementary Table 5). In total, we obtained 4,166,519 reads. After 97% OTU clustering and removing all non-bacterial and unassigned reads, a total of 3,080,204 reads remained. Samples were rarefied to 2,546 reads per sample and 10,525 OTUs overall were identified. Taxonomic diversity analysis revealed Proteobacteria, Firmicutes, Actinobacteria and Bacteroidetes to be the dominant bacterial taxa (Figure 8A), the same phyla identified on the lab-grown tomatoes. Alpha diversity analysis supported by the total number of observed species, and by the Shannon and Simpson diversity indices (Figure 8B) together with a pairwise comparisons using the Wilcoxon rank sum test showed significant differences in the number of OTUs between the hydroponic and the soil system (Supplementary Table 6).

**Figure 8.**
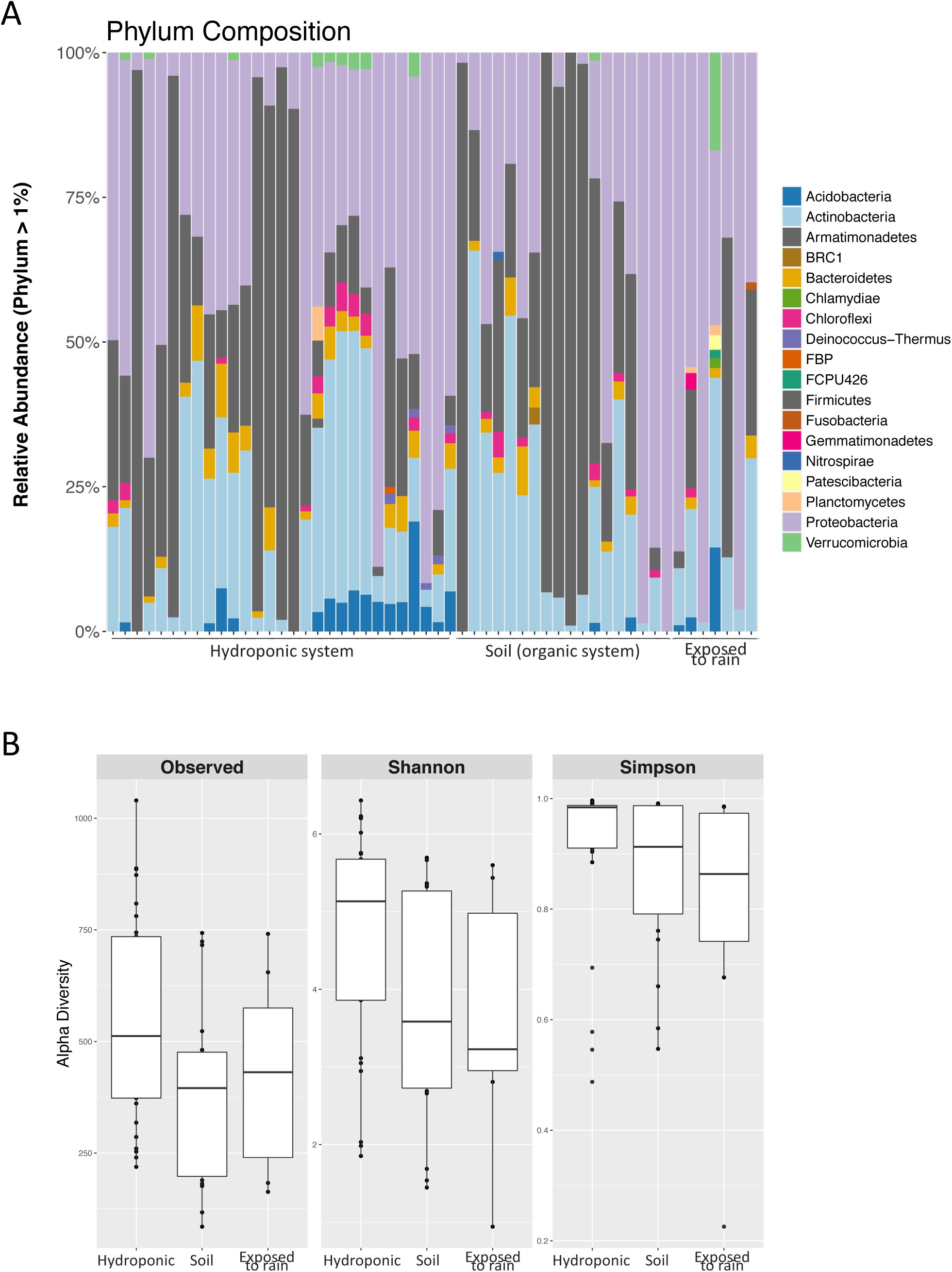
Taxonomic composition and Alpha diversity of phyllosphere microbiota of tomato plants grown hydroponically or in soil (both in a commercial greenhouse) and of tomato plants grown outside (on the roof of the Latham Hall research building). A) Relative abundance at the phylum level (only phyla with a RA above 1% are shown), B) Alpha diversity (observed OTUs, Shannon diversity index, and Simpson diversity index).

To identify any OTUs present at significantly higher relative abundance in tomato plants grown outside exposed to rain compared to the tomato plants grown in the greenhouse hydroponically in the absence of rain, DESeq2 was used again. Forty OTUs from the genera *Pandoraea*, *Curtobacterium*, *Massilia*, *Gemmatirosa*, *Kineococcus*, *Methylobacterium*, *Sphingomonas*, *Buchnera*, *Alloiococcus*, *Pseudomonas*, and *Streptococcus* were found (Figure 9A). Similarly, 58 OTUs from the genera *Pantoea*, *Pelomonas*, *Kineococcus*, *Methylobacterium*, *Massilia*, *Aureimonas*, *Buchnera*, *Alloiococcus*, *Pseudomonas*, *Streptococcus*, and *Sphingomonas* were of significantly higher relative abundance on the tomato plants grown outside exposed to rain compared to the tomato plants growing in soil never exposed to rain in the greenhouse (Figure 9B). Note that, as we hypothesized, OTUs in the genera *Massilia*, *Curtobacterium*, *Pseudomonas*, and *Pantoea*, were among the same genera as the OTUs identified to significantly increase in relative abundance in the phyllosphere of tomato plants treated with concentrated rain microbiota 7 days post inoculation. However, unexpectedly, the actual OTUs of these genera identified in this comparison were not the same as those identified in the controlled laboratory experiments (see discussion section for possible explanations).

**Figure 9.**
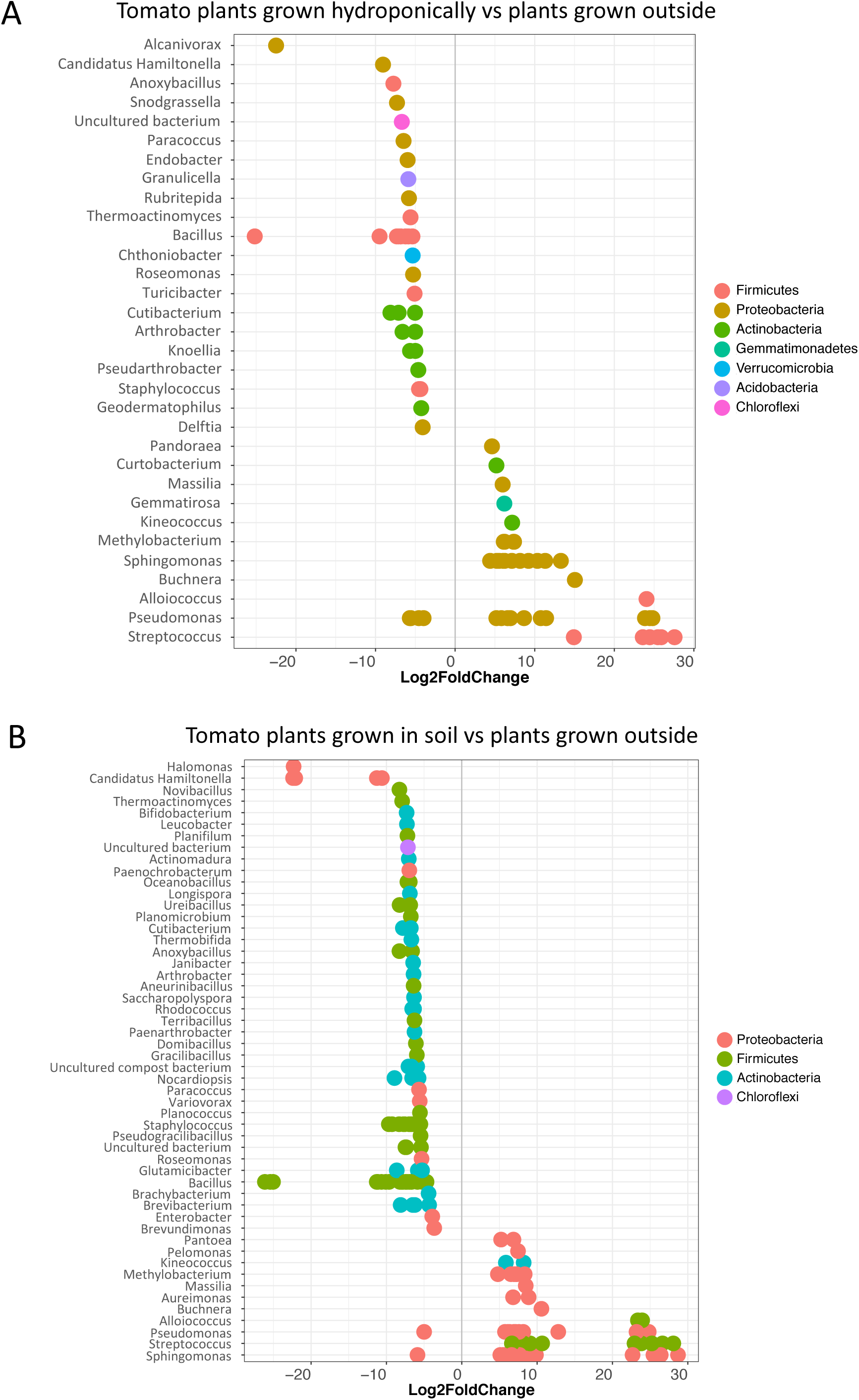
Differential abundance analysis at the level of Operational Taxonomic Unit (OTU) using DESeq2 (Love et al., 2014). The fold change is shown on the X axis and genera are listed on the Y axis. Each colored dot represents a separate OTU. A) Plants grown hydroponically in a greenhouse compared with plants grown on the roof of the Latham Hall research building, B) Plants grown in soil in a greenhouse compared with plants grown on the roof of the Latham Hall research building.

Finally, we performed another small experiment collecting rain on the same roof as in the previous experiments and exposing one set of tomato plants to all rain events while taking another set of plants inside before each rain event. From one rain event (June 10, 2019), phyllosphere communities from a single tomato plant exposed to rain (collected on June 21, 2019), and another tomato plant not exposed to rain (collected on the same date) were sequenced together with a sample of the corresponding rainfall using ONT’ MinION platform. A total of 194,591 long reads were analyzed using the ONT taxonomic classifier WIMP. Table 3 shows the most abundant species present in the three samples ranked by relative abundance on the tomato plant exposed to rain. Yet again, species of the genera *Massilia*, *Pantoea*, *Methylobacterium*, *Pseudomonas*, and *Janthinobacterium* (and some additional species outside of these genera) were present in the rain sample as well as in the tomato sample exposed to rain but absent (or almost absent) in the tomato sample not exposed to rain. While this experiment was too small for any statistical analysis, the result is consistent with the laboratory results as well as the comparison of tomatoes grown outside exposed to rain with those grown in a greenhouse not exposed to rain.

**Table 3.**
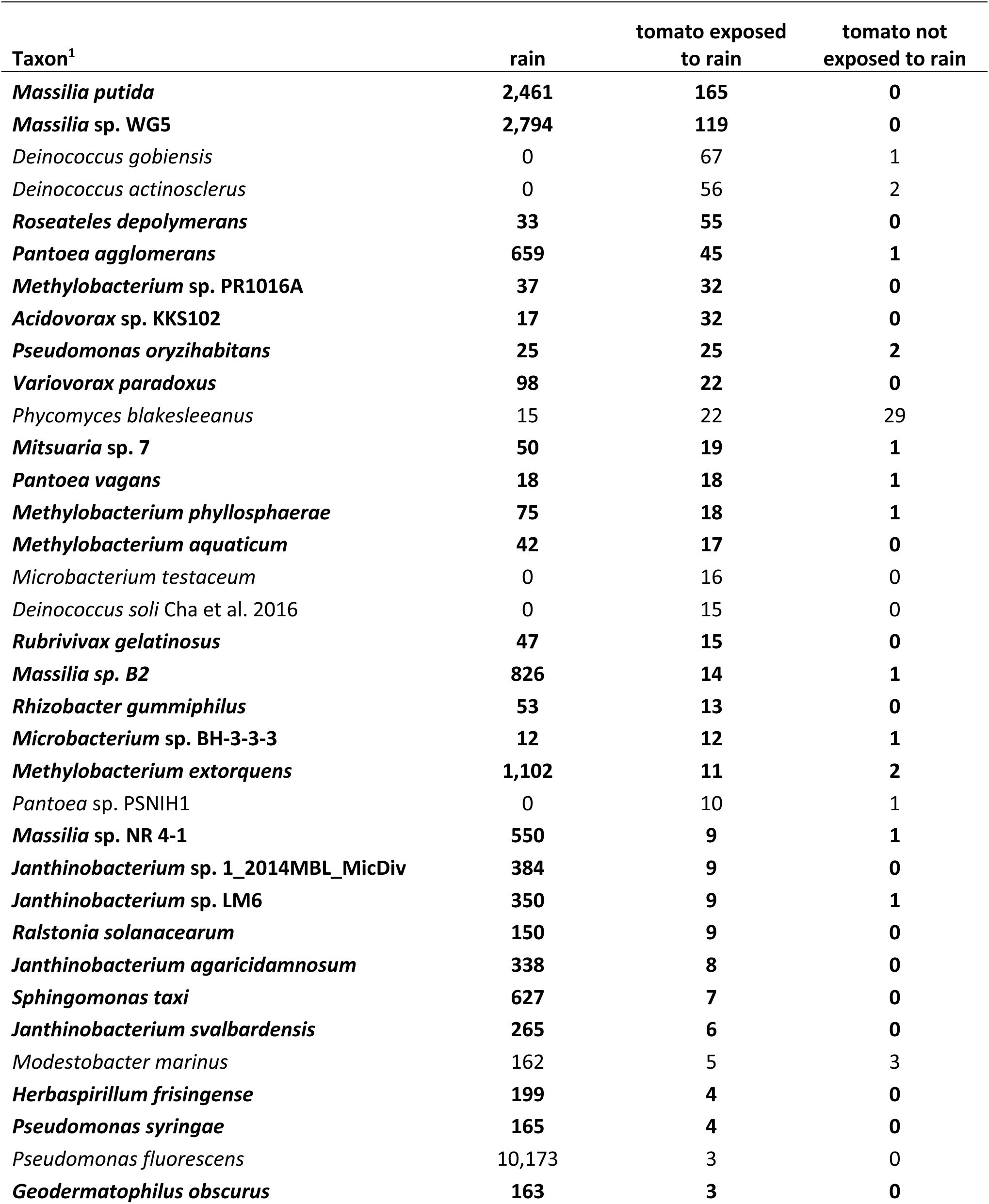
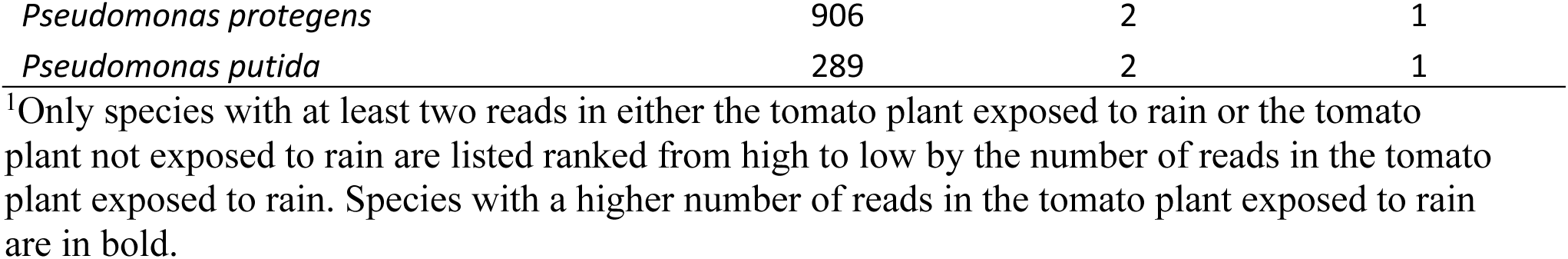
Taxonomic classification of ONT MinION reads using WIMP of a rain sample, a tomato plant exposed to rain, and a tomato plant not exposed to rain.

| **Taxon<sup>1</sup>** | **rain** | **tomato exposed**<br>**to rain** | **tomato not**<br>**exposed to rain** |
| --- | --- | --- | --- |
| ***Massilia putida*** | **2,461** | **165** | **0** |
| ***Massilia* sp. WG5** | **2,794** | **119** | **0** |
| *Deinococcus gobiensis* | 0 | 67 | 1 |
| *Deinococcus actinosclerus* | 0 | 56 | 2 |
| ***Roseateles depolymerans*** | **33** | **55** | **0** |
| ***Pantoea agglomerans*** | **659** | **45** | **1** |
| ***Methylobacterium* sp. PR1016A** | **37** | **32** | **0** |
| ***Acidovorax* sp. KKS102** | **17** | **32** | **0** |
| ***Pseudomonas oryzihabitans*** | **25** | **25** | **2** |
| ***Variovorax paradoxus*** | **98** | **22** | **0** |
| *Phycomyces blakesleeanus* | 15 | 22 | 29 |
| ***Mitsuaria* sp. 7** | **50** | **19** | **1** |
| ***Pantoea vagans*** | **18** | **18** | **1** |
| ***Methylobacterium phyllosphaerae*** | **75** | **18** | **1** |
| ***Methylobacterium aquaticum*** | **42** | **17** | **0** |
| *Microbacterium testaceum* | 0 | 16 | 0 |
| *Deinococcus soli* Cha et al. 2016 | 0 | 15 | 0 |
| ***Rubrivivax gelatinosus*** | **47** | **15** | **0** |
| ***Massilia* sp. B2** | **826** | **14** | **1** |
| ***Rhizobacter gummiphilus*** | **53** | **13** | **0** |
| ***Microbacterium* sp. BH-3-3-3** | **12** | **12** | **1** |
| ***Methylobacterium extorquens*** | **1,102** | **11** | **2** |
| *Pantoea* sp. PSNIH1 | 0 | 10 | 1 |
| ***Massilia* sp. NR 4-1** | **550** | **9** | **1** |
| ***Janthinobacterium* sp. 1_2014MBL_MicDiv** | **384** | **9** | **0** |
| ***Janthinobacterium* sp. LM6** | **350** | **9** | **1** |
| ***Ralstonia solanacearum*** | **150** | **9** | **0** |
| ***Janthinobacterium agaricidamnosum*** | **338** | **8** | **0** |
| ***Sphingomonas taxi*** | **627** | **7** | **0** |
| ***Janthinobacterium svalbardensis*** | **265** | **6** | **0** |
| *Modestobacter marinus* | 162 | 5 | 3 |
| ***Herbaspirillum frisingense*** | **199** | **4** | **0** |
| ***Pseudomonas syringae*** | **165** | **4** | **0** |
| *Pseudomonas fluorescens* | 10,173 | 3 | 0 |
| ***Geodermatophilus obscurus*** | **163** | **3** | **0** |
| *Pseudomonas protegens* | 906 | 2 | 1 |
| *Pseudomonas putida* | 289 | 2 | 1 |

<sup>1</sup>Only species with at least two reads in either the tomato plant exposed to rain or the tomato plant not exposed to rain are listed ranked from high to low by the number of reads in the tomato plant exposed to rain. Species with a higher number of reads in the tomato plant exposed to rain are in bold.

## DISCUSSION

While our general understanding of the plant microbiome has increased dramatically over the last few years, the basic question of where the bacteria that colonize and inhabit the phyllosphere originate from has remained unanswered. The main approach in trying to answer this question has been to make comparisons of the composition of the phyllosphere microbiome with the microbiomes of putative reservoirs (Maignien et al., 2014, Zarraonaindia et al., 2015, Ottesen et al., 2016, Wagner et al., 2016, Grady et al., 2019). To complement this approach, here we used controlled laboratory experiments.

We decided to focus on rain since previous studies provided evidence that at least the bacterial leaf pathogen, *Pseudomonas syringae*, may be disseminated by precipitation and efficiently colonize the plant phyllosphere (Monteil et al., 2016, Morris et al., 2008). Moreover, at least temporary shifts in the composition of phyllosphere communities after rain events were observed (Allard et al., 2020). In a first step, we found that tomato plants naturally exposed to rain outdoors, or simply sprayed with rainfall when grown in the lab, carried significantly higher bacterial populations sizes compared to lab-grown plants that had not been treated with rainfall or that were treated with sterilized rainfall. While plants grown outside may have also been colonized by airborne bacteria through dry-deposition or by soil-borne bacteria after being splashed during rainfall events, the observation that treating lab-grown plants with rain caused the bacterial population size to increase more than spraying lab-grown plants with sterilized rain was a strong indication that the observed increase in population size was due to the colonization and growth of rain-borne bacteria.

To dig deeper into the possibility that rain harbors bacteria that can effectively colonize and grow on tomato leaves, we decided to characterize the taxonomic composition of rainfall events and then determine if any of the identified members of the rain microbiota would increase in relative abundance on tomato plants treated with concentrated rain microbiota (100X-rain) but not on tomato plants treated with filter-sterilized rain or sterile water. We decided on using concentrated rain microbiota instead of rain since we had previously found that rain contained as few as 4 x 10^3^ CFU l^-1^ (Failor et al., 2017) and the maximum volume that we can spray on a tomato plant before water runs off the leaves is only 10 mL. Therefore, as few as 40 bacteria may get inoculated on an entire plant using rain as inoculum in a lab experiment, which we deemed not to be enough to lead to a bacterial population size that could be analyzed using 16S rRNA amplicon sequencing. We thus used 100X-rain concentrate, acknowledging that we artificially aided rain-borne bacteria in our experiment.

The most important observation from the characterization of the rainfall microbiota was that each collected rainfall event harbored a very different bacterial community. This is in line with other recent results on rainfall collected in the USA, Europe, and Asia (Aho et al., 2020, Cáliz et al., 2018, Woo & Yamamoto, 2020) that showed that the taxonomic composition of microbiota in rain changes with origin of air masses and season. This meant for our experiment that we could not expect to find the same taxa to colonize and grow on tomatoes in each inoculation experiment.

It was also important to use appropriate controls when inoculating lab-grown tomato plants with 100X-rain. Importantly, tomato plants were not grown in sterile conditions. Therefore, they already carried microbiota at the time of inoculation and simply spraying these plants with water and incubating them at high humidity (as we did to favor plant colonization) could be expected to lead to changes in relative abundance of the pre-existing microbiota. Moreover, rain contains nutrients and bacteriophages that are not removed by filtering and that will also affect pre-existing microbiota. Therefore, we used both, filter-sterilized rain (including nutrients and possibly bacteriophages but no bacteria) and autoclaved double-distilled water (expected to contain neither nutrients nor bacteriophages nor bacteria) as controls.

Comparison of rain microbiota with phyllosphere microbiota on day-0 and day-7 for the three different treatments yielded some results confirming our hypothesis that rain contains effective colonizers of tomato leaves while other results were ambiguous. Firstly, the observed drop in alpha diversity for phyllosphere microbiota from day-0 to day-7 after treatment with 100X-rain is in agreement with the expected increase in relative abundance of some effective tomato leaf colonizers accompanied by a decrease in relative abundance of many rain-borne bacteria that are not adapted to tomato leaves and that are thus not effective tomato colonizers. The observed decrease in alpha diversity from day-0 to day-7 for the sterile rain treatment may have been due to growth of some pre-existing tomato phyllosphere members that thrived under the high humidity conditions applied for two days after inoculation and the nutrients added with the rain water, which helped them outcompete other members of the pre-existing microbiota. It is also possible that bacteriophages present in the filter-sterilized rain reduced the relative abundance of some phyllosphere members to below the detection limit.

In regard to beta diversity, we had expected the composition of the day 0 microbiota to be similar to each other because tomato plants of the same cultivar were grown in autoclaved soil in relatively stable laboratory conditions. Therefore, it was surprising to find phyllosphere samples to differ more from each other (even on day-0 after sterile water treatments) compared to the differences among rain samples. This was the case for weighted UniFrac distances, unweighted UniFrac distances, and the Bray-Curtis dissimilarity static. A possible explanation is that the composition of the phyllosphere microbiota of our lab-grown tomatoes was determined by stochastic processes because of the low concentration of plant-associated bacteria present in the indoor air and the autoclaved soil that was used for growing.

Another unexpected result was that beta diversity significantly changed from day-0 to day-7 for all three treatments. Although the high humidity maintained for two days after inoculation was expected to have some effect on the phyllosphere community, we still expected 100X-rain to have a stronger effect on beta diversity than sterile water. Similarly unexpected, the taxonomic composition at the phylum, class, and genus level on day-0 and day-7 did not reveal any consistent change unique to the treatment with 100X-rain compared to filter-sterilized rain or sterile water. Taken together, these results suggested that, if they existed, tomato leaf colonizers present in rain were individual species and changes in abundance of these individual species were not evident from the overall comparison of beta diversity or taxonomic composition at higher taxonomic ranks.

Therefore, we decided to look at changes at the OTU level. To do this, we used DESeq2 (Love et al., 2014) a tool originally developed to identify significant changes in gene expression in RNA-seq experiments, which have similar challenges to OTU tables, and which has been shown to be effective for smaller OTU datasets like ours (Weiss et al., 2017). We made several comparisons. Most importantly, not a single OTU significantly increased from day 0 to day 7 after treatment with filter-sterilized rain or sterile water but 104 OTUs increased significantly after treatment with 100X-rain. Since none of these OTUs significantly increased from day-0 to day-7 after treatment with filter-sterilized rain or sterile water, they were most likely rain-borne.

The 104 OTUs belong to 10 genera with one of them being the genus *Pantoea*. The DeSeq2 analysis identified 18 OTUs of this genus that significantly increased in relative abundance from day 0 to day 7 after being treated with 100X-rain. Importantly, *Pantoea* OTUs were also present in all rain samples (including the one sequenced with ONT’s MinION). Moreover, the species *P. agglomerans* and *P. vagans* were identified as the most abundant species in several of the tomato phyllosphere day 7 samples after being treated with 100X-rain based on metagenomic sequencing. Also, two *Pantoea* OTUs were present in significantly higher relative abundance in tomatoes grown outside compared to tomato grown organically inside a commercial greenhouse, and *P. agglomerans* and *P. vagans* were both identified in the rain metagenome as well as in the metagenome of a tomato plant exposed to rain but not in a tomato plant not exposed to rain using ONT MinION sequencing. Moreover, the genus *Pantoea* is well known to include plant pathogenic bacteria and beneficial plant-associated bacteria (Coutnho & Venter, 2009, Walterson & Stavrinides, 2015, Mechan Llontop et al., 2020), we previously identified 192 *Pantoea* isolates in a culture-dependent study of precipitation samples (Failor et al., 2017), and *Pantoea* species were recently identified both in rainfall before and after falling through a forest canopy, with higher relative abundance in the throughfall samples (Ladin et al., 2021). Therefore, based on the results obtained here and data in previous literature, members of the genus *Pantoea* are likely phyllosphere inhabitants that originate from rainfall.

Another rain-borne genus that includes species that appear to successfully colonize tomato plants is *Massilia*. Members of this genus were found in all rain samples (those analyzed by 16S rRNA amplicon sequencing and the one sequenced with ONT’s Minion). Twenty-seven OTUs of this genus significantly increased between day-0 phyllosphere samples and day-7 samples for 100X-rain treated plants. Two *Massilia* species were also found among the species with the highest relative abundance in the metagenomic sequences of the 100X-treated tomato samples on day 7. One *Massilia* OTU each was more abundant in tomatoes grown outdoors compared to hydroponically or organically grown tomatoes in the commercial greenhouse, respectively. Four *Massilia* species were found in the rain sample and tomato sample grown outdoors but not in the tomato plant not exposed to rain when using ONT MinION sequencing. As with *Pantoea*, *Massilia* species were cultured out of precipitation by us previously (Failor et al., 2017) and were recently found in rain and rain that had fallen through a forest canopy (Ladin et al., 2021). Finally, OTUs and named species belonging to the genus *Massilia* have been found in plants, soil, and even extreme environments (Ofek et al., 2012, Rastogi et al., 2012, Bodenhausen et al., 2013, Purahong et al., 2018, Singh et al., 2019, Holochová et al., 2020). Therefore, members of the genus *Massilia* may cycle through multiple environments and some of them may be transported by rain to leaf surfaces where they colonize the phyllosphere.

Other rain-borne genera likely to colonize the tomato phyllosphere based on our data include *Janthinobacterium*, which, as the genus *Massilia*, is a member of the Burkholderiaceae family. Four *Janthinobacterium* species were also found in rain and in the rain-exposed tomato plant but not in the tomato plant protected from rain in the ONT MinION experiment. However, *Janthinobacterium* was not found at significantly higher relative abundance in tomato plants grown outside compared to greenhouse-grown tomatoes. Its inconsistent presence in rain may explain this result.

Finally, we found evidence for OTUs and named species of the genus *Pseudomonas* to colonize tomato plants. Unexpectedly though, we did not find a single member of the *Pseudomonas syringae* species complex (*P. syringae sensu lato*), which includes many plant pathogenic and commensal lineages (Vinatzer et al., 2016, Monteil et al., 2016). We do not have any good explanation why we neither found *P. syringae* at relatively high abundance in the analyzed rain samples nor in our phyllosphere samples treated with 100X-rain although we had previously cultured *P. syringae* from rain (Failor et al., 2017) and from plants in our geographic area (Clarke et al., 2010). We conclude that while *P. syringae* pathogens are present in rain and disseminated by precipitation, they may not be a major component of precipitation microbiota, at least not in the geographic area where the experiments here described were performed.

Importantly though, while our data and the literature suggest that members of some common phyllosphere genera are rain-borne, the fact that we found different enrichment of OTUs from these genera across experimental conditions, lab versus greenhouse, precluded the identification of likely rain-borne tomato phyllosphere colonizers at the species level. One possible explanation for this is that each rain event harbored such different taxa and this increased variation coupled with low replication meant that OTUs with higher relative abundance on tomatoes grown outside were simply missed in the lab experiments. Conversely, the small number of analyzed tomato plants grown outside exposed to rain made it difficult to find significant differences compared to the tomatoes grown inside not exposed to rain. Finally, laboratory conditions may not have allowed some of the OTUs found outside to effectively grow on tomato plants inside the laboratory. On the other hand, artificial light and almost constant temperature and humidity may have favored tomato colonization of related, but different, species compared to the most effective colonizers of tomato plants grown outside, where plants are exposed to natural sunlight, including UV radiation and dramatic temperature and humidity changes. These differences in environmental conditions may explain why members of the genera *Sphingomonas* and *Methylobacterium* were consistently found in rain and in significantly higher abundance on tomato plants grown outside than in greenhouse-grown tomatoes but OTUs of these genera did not significantly increase in our controlled laboratory experiments after 100X-rain treatments. Finally, the use of concentrated rain microbiota instead of rain may have led to increased competition between rain-borne bacteria and suppressed the colonization efficiency and growth of some while favoring the growth of others.

To follow up on the results reported here and to gain further insight into the relative importance of seeds, soil, the atmosphere, and precipitation as reservoirs of phyllosphere microbiota, we propose to expand the kind of controlled experiments we performed here but growing tomatoes outdoors from either sterilized seeds or non-sterilized seeds, either not limiting exposure to precipitation or limiting exposure to precipitation (for example, through the use of mobile rain-out shelters), and growing plants in native versus sterile soil. Moreover, strain-level metagenomics (Olm et al., 2021) of all reservoir microbiota and phyllosphere microbiota could provide the necessary strain-level resolution to identify which strains from which reservoirs are the most important colonizers of the phyllosphere.

## DATA DEPOSITION

Sequences and metadata are being deposited at NCBI under BioProject DPRJNA719680. All read processing steps, bioinformatic workflows, and scripts used in this research are available on GitHub (https://github.com/marcoeml/VinatzerLab-Mechan-rain-phyllosphere-microbiota-2020).

---

**Supplementary Figure 1.**
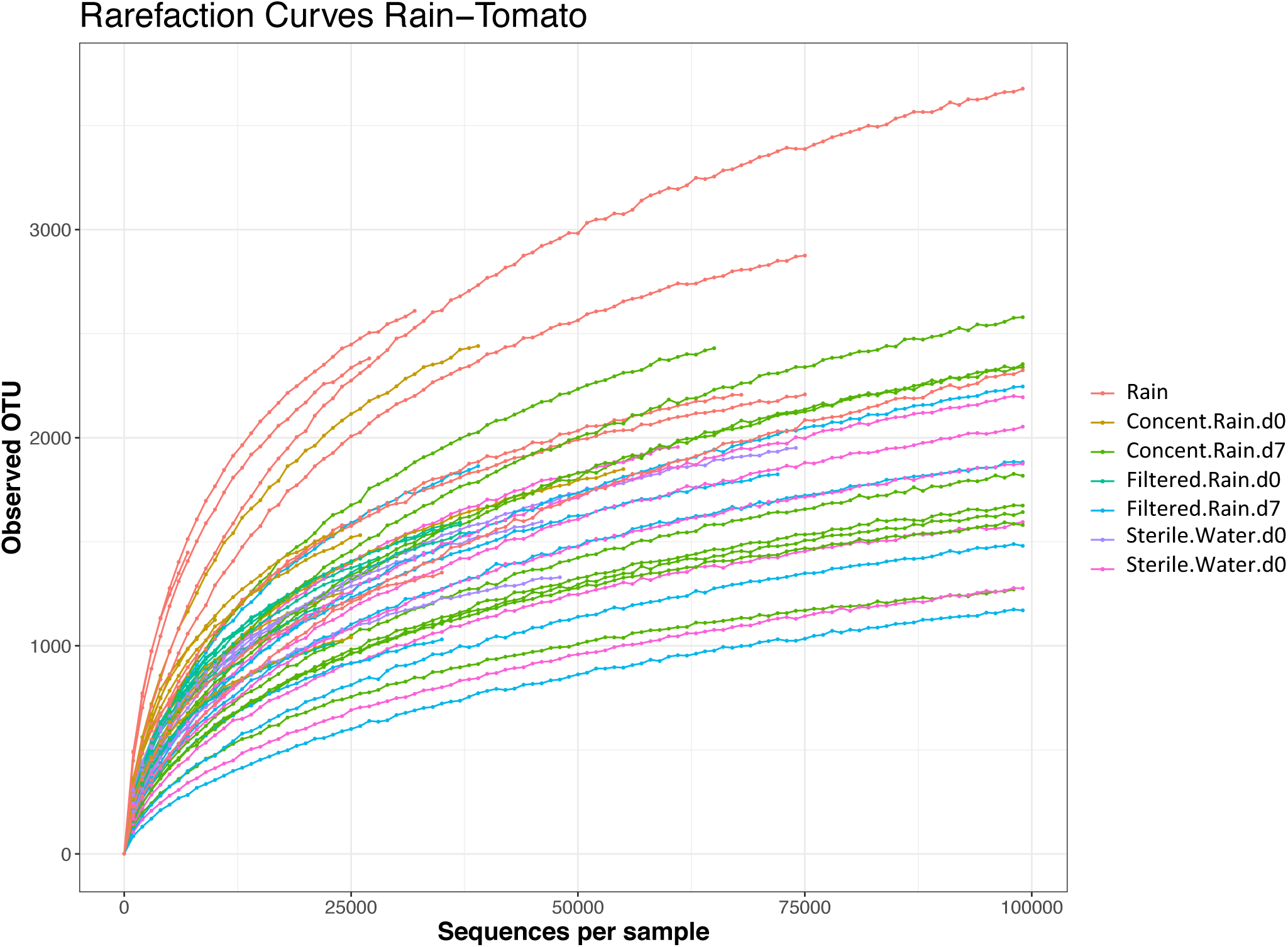
Rarefaction curves of rain microbiota and tomato phyllosphere microbiota treated with 100X-rain, filtered rain, or sterile water on day-0 and day-7.

**Supplementary Figure 2.**
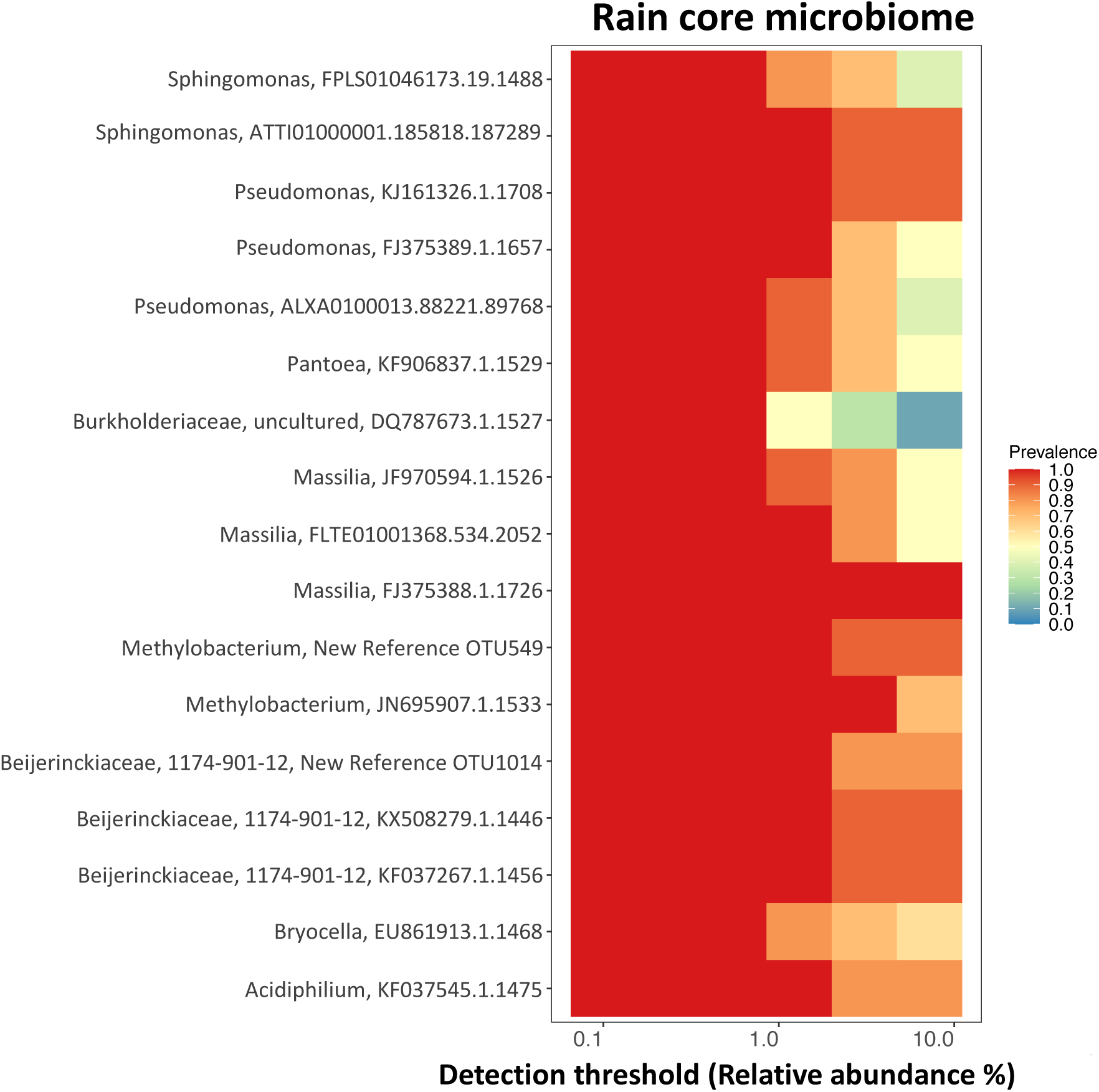
The core rain microbiome. Rain-associated OTUs at a detection threshold of 0.1% and 100% as a prevalence threshold.

**Supplementary Figure 3.**
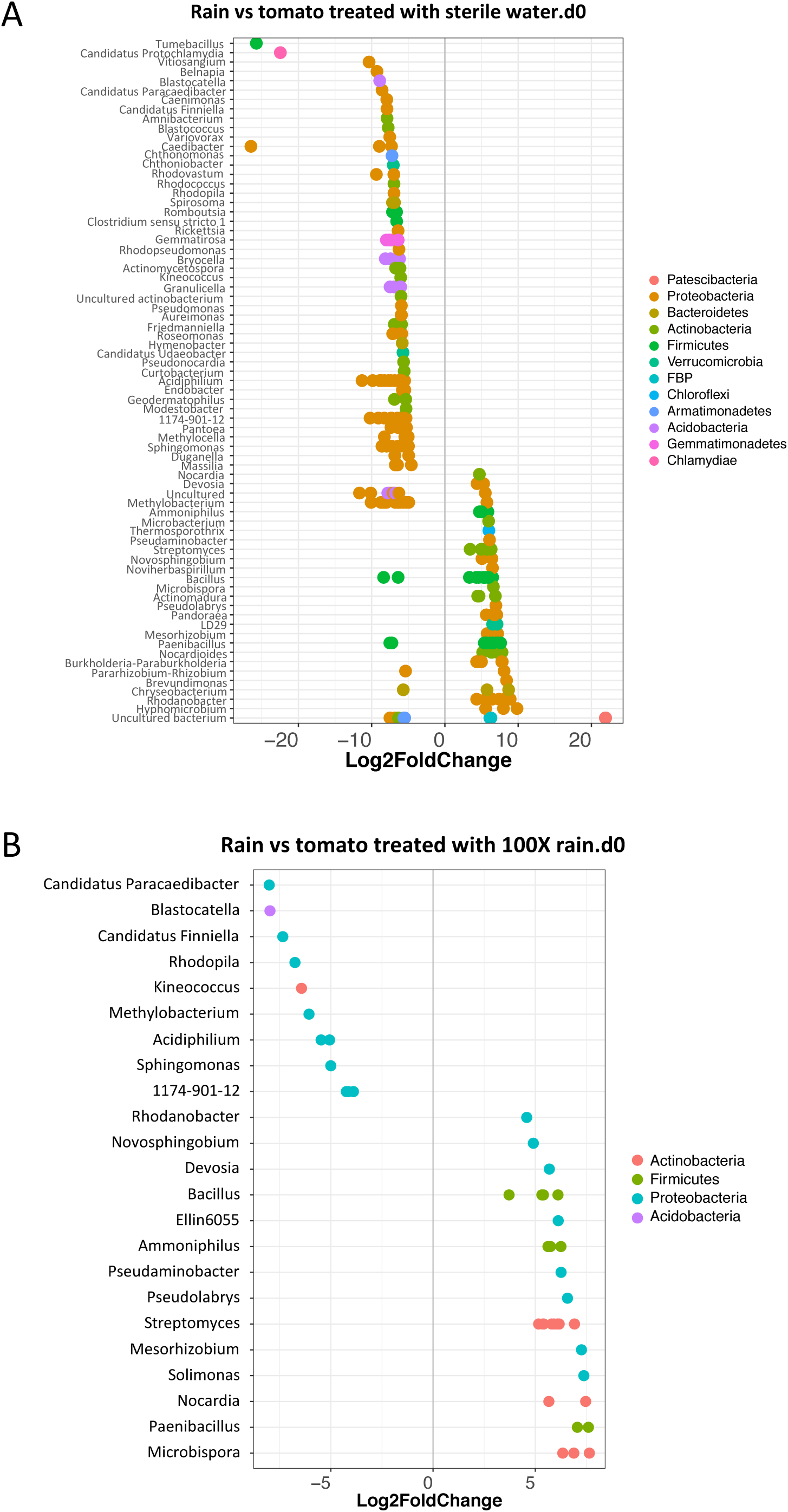
Differential abundance analysis at the OTU level usingDESeq2 (Love et al., 2014). A) Comparison of rain microbiota with the tomato phyllosphere treated with sterile water at day-0 during the same experiment, B) Comparison of rain microbiota with the tomato phyllosphere microbiota on the day there were treated with 100X-rain (day-0).

